# Lysosome-dependent nutrient scavenging underlies stress adaptation during epithelial-to-mesenchymal transition

**DOI:** 10.1101/2025.09.22.677807

**Authors:** Michal J. Nagiec, Paola M. Cavaliere, Friederike Dundar, Derrick M. Spencer, Nikolaos Koundouros, Joana B. Nunes, Sejeong Shin, Sang-Oh Yoon, Julie Han, Andre Chavez, Olivia M. Kester, Rabia Khan, Paul Zumbo, Eric E. Gardner, Bobak Parang, Anders P. Mutvei, John M. Asara, Steven P. Gygi, Doron Betel, Noah Dephoure, John Blenis

## Abstract

Metastatic cancer cells invade tissue, overcome nutrient stress, and survive transit to distant sites. Many of the mechanisms that support these processes are incompatible with proliferation. This study defines cellular transition states in breast epithelial cells undergoing epithelial-mesenchymal transition (EMT) driven by ERK2 and TGF-β signaling. EMT triggers robust endolysosomal system upregulation and metabolic adaptations that balance proliferative and invasive states. Surprisingly, invasive cells rely on scavenging via lysosomes and macropinocytosis to acquire amino acids, rather than plasma membrane transport, even in nutrient-rich conditions. Macropinocytosis increases intracellular amino acid storage, promoting survival during amino acid deprivation. This metabolic shift depends on c-MYC downregulation, an early EMT event. Reintroducing c-MYC suppresses the metabolic switch, endolysosomal induction, macropinocytosis, and the proliferation-to-migration transition. These findings reveal how cells dynamically balance proliferation and invasion, offering insights into transition states difficult to capture in models of breast cancer metastasis.

---

Metastasis is the leading cause of cancer mortality and the product of a selection process that generates highly resilient cells. Cancer cells that reach this advanced stage become invasive and more resistant to a broad range of stresses allowing for spread to distant organs. These include nutrient and mechanical stress, free radicals, hypoxia and genotoxic stress generated by nontargeted therapies^1, 2^. Common to metastatic cells is the reactivation of gene transcripts associated with the developmental program, epithelial-to-mesenchymal transition (EMT). In tissue development and repair, the physiologic EMT process is transient and tightly controlled, where cells are restored to their differentiated state. Pathologic EMT activation in tumors, however, is dysregulated, often incomplete and sustained, conferring cells with the capacity to invade distant sites while mitigating associated stress. Identifying targetable vulnerabilities of pathologic EMT is fundamental to devising strategies for reducing cancer mortality.

The EMT state is controlled by diverse signaling networks including TGF-β, Ras, Akt, and β-catenin^3, 4^. Despite the heterogeneity of signals promoting EMT, they frequently converge at the activation of ERK1/2 in various cancers^5–8^. Our studies of a gain-of-function ERK2^D319N^ mutant have identified key signaling, transcriptional, and chromatin remodeling events downstream of ERK2 required for activation of an EMT program that results in enhanced invasive capacity^9–11^, while other studies have highlighted the relevance of ERK2 in driving adaptive signaling and increasing metastatic burden by driving EMT responses^12, 13^. Importantly, nascent metastatic tumor cells may need EMT signaling not just to invade but also to adapt to changing nutrient availability encountered during metastatic transit^14^.

Lysosomes play critical roles in mitigating nutrient stress and breaching barriers to promote tumor dissemination^15, 16^. Cancer cells facing a shortage of nutrients or increased energetic demand, utilize extracellular scavenging and intracellular recycling programs to catabolize macromolecules^17^. Both intracellular and extracellular scavenging programs require fusion with lysosomes to catabolize engulfed macromolecules. In various cancers, including breast cancer, adaptation to nutrient limited conditions by tumor cells has been associated with more aggressive disease^18, 19^. Notably, mutant Ras driven pancreatic ductal adenocarcinoma (PDAC) cells that are scavenging competent exhibit EMT phenotypes after prolonged amino acid starvation suggesting a potential coordination between nutrient stress adaptation and signaling associated with metastatsis^20, 21^. However, understanding how EMT signaling and nutrient scavenging converge in normal and tumor cells is likely to reveal mechanisms critical to metastatic tumor development.

An emerging strategy in combating advanced stage tumors is the identification of tumor cell dependencies that increase their adaptability and survival^14, 22^. We set out to identify cellular programs engaged during EMT that facilitate the acquisition of phenotypic plasticity and support invasive behavior. To capture and evaluate these events we conducted time resolved multi-omic analysis of ERK2-driven EMT. We find that EMT engages in reciprocal signaling between ERK2 and TGF-β. This signaling is associated with a pronounced shift in cell metabolism as cells acquire invasive capacity. The metabolic shift is accompanied by an increase in the endolysosomal compartment and increased rates of nutrient scavenging and is marked by suppression of glutamine and essential amino acid (EAA) transporters at the plasma membrane (PM) and increased expression of the lysosomal glutamine transporter SNAT7. The elevated flux through the endolysosomal system enhanced cellular amino acid stores, allowing survival during prolonged nutrient stress. Cells engaged in EMT become sensitive to inhibition of scavenging or lysosome inhibitors which also block invasive capacity of triple negative breast cancer cells. We find that suppression of the proto-oncogene c-MYC during EMT is critical for this transition and restored c-MYC protein expression results in a robust repression of the endolysosomal response and reversal of the metabolic shift. Together our data highlight a network of critical nodes engaged in the transition from a proliferative to invasive state and cement the importance of amino acid sensing and acquisition through endolysosomes.

## Results

### A gain of function ERK2 mutant promotes EMT and altered cell metabolism

Tumor associated EMT is characterized by a trade-off between proliferation and migration, and increased resilience to tumor associated stress. We set out to capture dependencies associated with epithelial plasticity by characterizing cellular dynamics of EMT. We measured temporal changes in mRNA, protein, and metabolites after inducible expression of a gain of function ERK2 mutant (Rat D319N corresponds to Human D321N) in non-tumorigenic MCF10A breast epithelial cells (Fig. 1a). ERK2^D319N^ expression drives a EMT response with strong fidelity that can be monitored temporally for the assessment of early (24h), intermediate (3-5 days), and late (7-9 days) phases of the transition^9–11^ (Fig. 1b and Extended Data Fig. 1a). Although, no change to cell morphology is observed after 24h, we noted an increase in the appearance of lamellipodial structures and cell spreading three days after ERK2^D319N^ induction along with increased vacuolization indicative of altered vesicular trafficking (Fig. 1b). Mapping tumor-associated MAPK1 (encoding ERK2) gene mutations onto the structure of the common docking (CD) site identified two additional residues found mutated in breast and various squamous cell carcinoma tumors, including head and neck^23^ (Extended Data Fig. 1b,c). We found that expression of these mutants phenocopied ERK2^D319N^, including loss of E-cadherin and induction of fibronectin and stabilization of the EMT transcription factor ZEB1, along with loss of signaling to the downstream kinase p90 RSK (Extended Data Fig. 1d). Cells expressing vector or ERK2^Y261A^, harnessing a mutation in the DEF-binding pocket required for EMT^9^, maintained an epithelial phenotype over the course of our experiment and served as controls, while induction of ERK2^WT^ and ERK2^D319N^ expression resulted in morphological and protein changes associated with EMT (Fig. 1b and Extended Data Fig. 1a). Dynamic profiles of transcript and protein expression in response to ERK2^D319N^ induction were clustered temporally (Fig. 1c,d). Enrichment analysis confirmed cells engaged in EMT exhibited early suppression of E2F, MYC and cell cycle associated gene sets with a steadily increasing induction of gene sets associated with EMT, TNF-α, TGF-β, and hypoxia (Fig. 1e,f). Among the hallmark gene sets ERK2^D319N^ expressing cells showed notable overlap with previously described TGF-β response associated with breast tumor lung metastasis^24^ (Extended Data Fig. 1e) and patterns observed among breast tumor cell lines after metastatic colonization of distinct organs^25^ (Extended Data Fig. 1f,g). Interestingly, our temporal analysis revealed differential expression of EMT transcription factors with Slug expression observed by day 1, ZEB1 by day 3 followed by ZEB2 and Twist (Extended Data Fig. 1h). Together these data highlight that ERK2^D319N^ induction elicits a cellular reconfiguration that favors cell migration, while limiting proliferation, and corresponds to gene expression changes associated with metastatic spread.

**Fig. 1.**
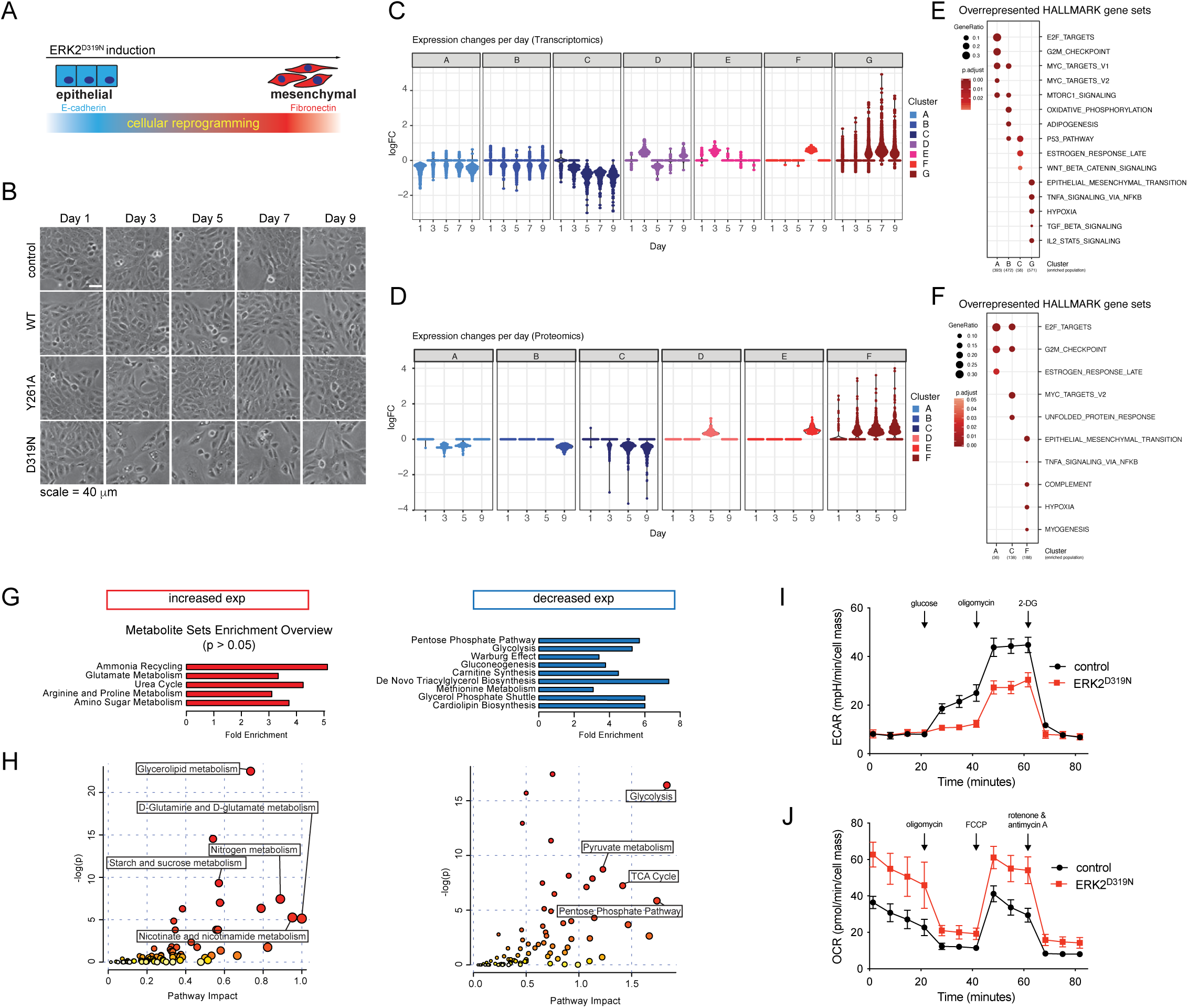
A gain of function ERK2 mutant promotes EMT and altered cell metabolism. **a**, Schematic of an inducible ERK2^D319N^ driven EMT model. **b**, Brightfield images of MCF10A cells expressing vector control, ERK2 wild type (WT) or mutants (Y261A, D319N) in response to 250 ng/ml doxycycline over 9 days. **c**, Groups of differentially expressed gene transcripts collected from cells grown as in **b**, plots show clustering analyses using k-nearest neighbors (kNN) of statistically significant expression changes. **d**, As in **c,** but for differentially expressed proteins. **e**, Enrichment analysis of clusters shown in **c**. **f**, Enrichment analysis of clusters shown in **d**. **g**, Enrichment analysis of differentially expressed metabolites. **h**, Joint transcript and metabolite pathway analysis of those with increased or decreased expression. **i**, Glycolysis stress test profile indicating changes to the extracellular acidification rate (ECAR) of control and ERK2^D319N^ expressing cells on day 3, (n ≥ 10). **j**, Mitochondrial stress test profile indicating changes to the oxygen consumption rate (OCR) of control and ERK2^D319N^ expressing cells on day 3, (n ≥ 10).

To capture metabolic changes that accompany this transition we collected metabolites from control and ERK2^D319N^ expressing cells. Pathway enrichment analysis of differentially expressed metabolites revealed a decrease in glycolysis and pentose phosphate pathway associated metabolites and an increase in metabolites associated with ammonia recycling, glutamate and amino sugar metabolism as epithelial cells transition to a more mesenchymal state (Fig. 1g). The observed shifts in carbon and nitrogen metabolism were further supported by joint transcriptomic and metabolomic pathway analysis. The decrease in glucose metabolites was accompanied by a decrease in the transcripts of glycolytic enzymes (Fig. 1h). To measure glycolysis, we monitored extracellular acidification using a Seahorse analyzer and found ERK2^D319N^ expressing cells yielded a marginal response to glucose and exhibited roughly half of the glycolytic capacity of control cells (Fig. 1i). Metabolic plasticity is associated with the ability of cells to compensate shifts in glycolysis with mitochondrial respiration^26, 27^. To explore this relationship, we measured mitochondrial respiration and found an inverse response in ERK2^D319N^ expressing cells, where the rate of basal and maximal mitochondrial respiration was roughly doubled in cells undergoing EMT (Fig. 1j). These data suggest a metabolic shift away from glycolysis in highly proliferative cells toward mitochondrial respiration in invasive cells.

### ERK2^D319N^ engages TGF-β signaling to drive EMT

The TGF-β pathway is a principal driver of EMT programs in tissue repair and cancer, where it cooperates with the RAS-ERK pathway to induce downstream effects^28^, and whose inhibition diminishes the TGF-β response^29^ (Extended Data Fig. 2a). ERK2^D319N^ expressing cells exhibit enriched expression of the TGF-β hallmark gene set (Fig. 1e), and SERPINE1/PAI1, a TGF-β activated gene/protein, is increased 24 hours after ERK2^D319N^ induction (Fig. 2a and Extended Data Fig. 1e), suggesting ERK2 activates EMT in cooperation with TGF-β signaling. To determine if ERK2^D319N^ driven EMT requires TGF-β signaling, cells were treated with vactosertib, an inhibitor of TGF-β receptor kinase type I (TGFBRI/ALK5). Vactosertib blocked EMT progression assessed by a suppression of morphological changes (Fig. 2b), gene expression patterns of EMT markers (Fig. 2c) and invasion (Fig. 2d). Furthermore, knockdown of SMAD2, SMAD3, or SMAD4, key TGF-β effectors, significantly inhibited ERK2^D319N^-driven invasion (Extended Data Fig. 2b,c). We also observed increased SMAD3 phosphorylation on the carboxy tail when SMAD3 was overexpressed in ERK2^D319N^ expressing cells without a notable change in SMAD2 phosphorylation (Extended Data Fig. 2d). SMAD3 overexpressing cells exhibited a stronger EMT phenotype under TGF-β treatment with dramatically reduced E-cadherin and elevated fibronectin, while the SMAD3 C-terminal truncation mutant acted as a dominant negative (Extended Data Fig. 2e-g). Notably, expression of a SMAD3 mutant that blocks receptor mediated phosphorylation suppressed PAI1 and fibronectin induction, restored E-cadherin expression and reversed the TGF-β mediated increase in ERK2 phosphorylation highlighting SMAD3 as a critical node in ERK2 and TGF-β signal coordination (Extended Data Fig. 2e). Together these data indicate ERK2-driven EMT engages TGF-β signaling to promote invasion and reciprocally TGF-β signaling requires ERK2 function.

**Fig. 2.**
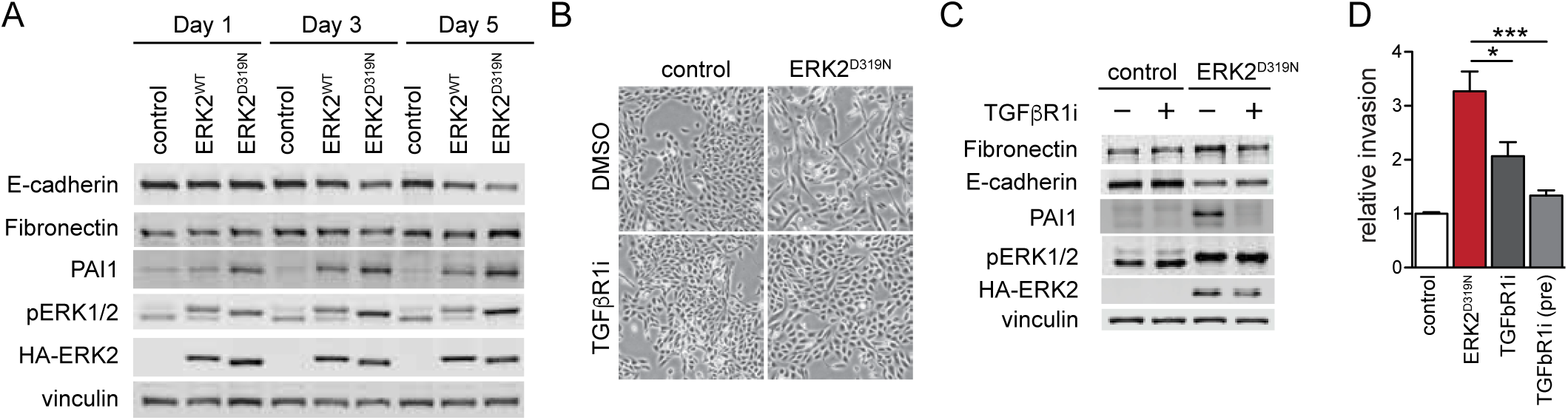
ERK2^D319N^ engages TGF-β signaling to drive EMT. **a**, Immunoblot of MCF10A cells expressing vector control, ERK2^WT^ and ERK2^D319N^ for 1, 3 and 5 days induced with 250 ng/ml doxycycline. **b,** Brightfield images of MCF10A cells after five days induction with 250 ng/ml doxycycline and treatment with 5 μM TGF-β receptor inhibitor (vactosertib). **c**, Immunoblots corresponding to **b**. **d**, Matrigel invasion assay of cells after 7 days of doxycycline. Cells treated with 5 μM vactosertib when seeded in Boyden chamber or (pre) cultured two days in 5 μM vactosertib prior to chamber seeding (n=3). Values are expressed as mean ± SEM (*p < 0.05, **p < 0.01, ***p < 0.001).

### EMT signaling drives lysosomal induction

The convergence of ERK2 and TGF-β signaling in EMT prompted further investigation into how these pathways cooperate to drive the transition to an invasive state. Quantitative proteomics of ERK2^D319N^-cells and of cells undergoing TGF-β-driven EMT revealed a remarkably high concordance between the two systems in both the suppression and induction of differentially expressed proteins (Extended Data Fig. 3a,b). We observed increases among EMT-associated proteins and decreases among proliferation-associated proteins in both systems (Fig. 3a). Especially striking was the increase in lysosomal proteins that was common to both EMT models, suggesting a potential cellular dependency of EMT on mobilization of lysosomal functions (Fig. 3b and Extended Data Fig. 3c). In the ERK2^D319N^ model and in the presence of nutrient-rich conditions, lysosomal proteins increased on day 3 and were sustained as cells progressed through EMT (Fig. 3c). The increase in lysosomal proteins coincided with an appearance of vacuoles three days after ERK2^D319N^ induction (Fig. 1b) and elevated expression of active form cathepsin proteases found in the lysosomal lumen (Fig. 3d).

**Fig. 3.**
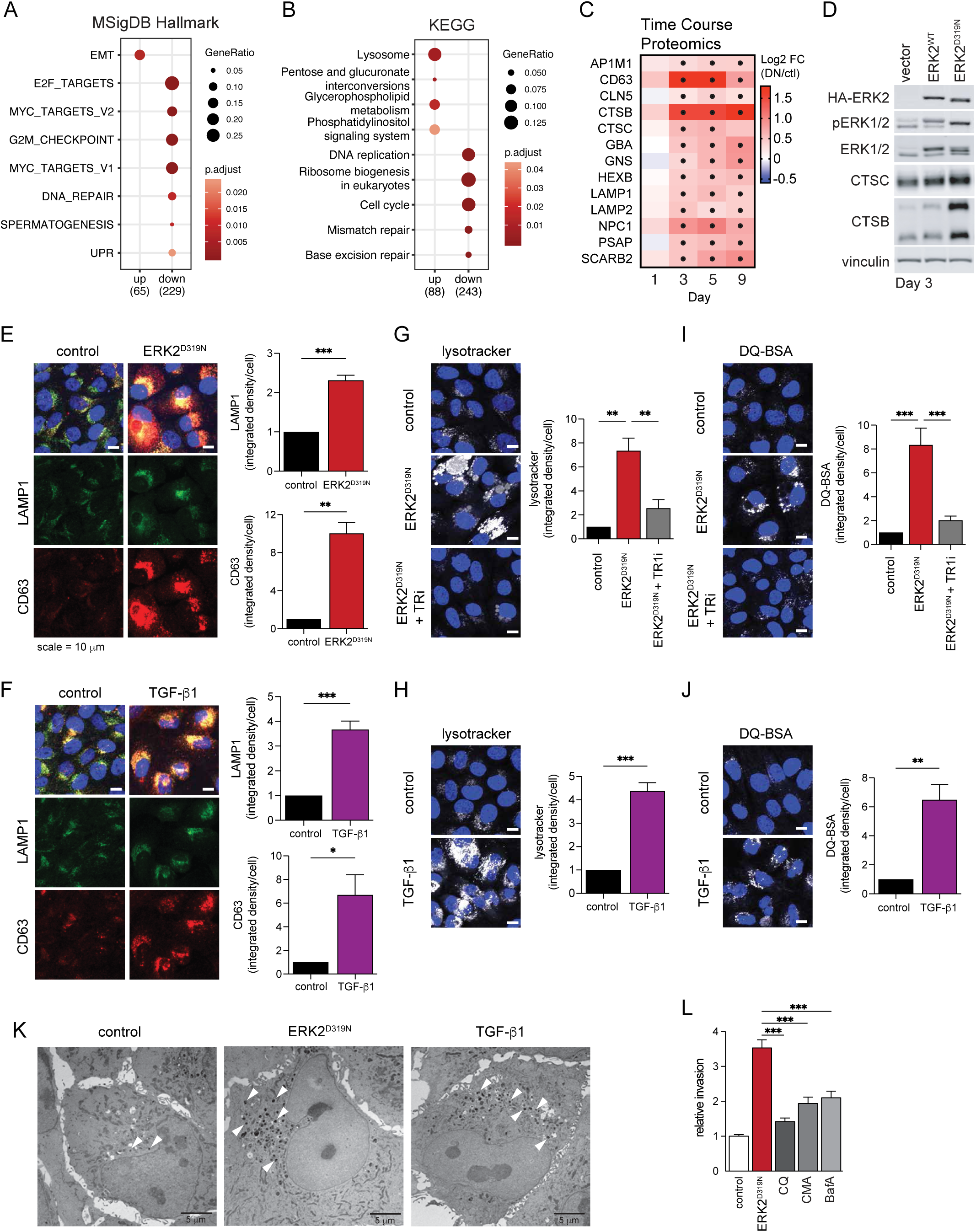
EMT signaling drives lysosomal induction. **a**, Enrichment analysis of up and down-regulated proteins in the two models of EMT using MSigDB Hallmark terms. **b**, As in **a**, but using KEGG terms. **c**, Lysosome associated protein expression changes after ERK2^D319N^ induction. Values represent fold change relative to the starting time point (n = 4). **d**, Immunoblot of MCF10A cells expressing ERK2^WT^ and ERK2^D319N^ cells after 3 days. **e**, LAMP1 and CD63 immunofluorescence of MCF10A ERK2^D319N^ expressing cells. **f**, As in **e**, cells treated with 5 ng/ml TGF-β1 (**e** and **f** each graph, n = 3). **g**, Lysotracker signal in MCF10A ERK2^D319N^ expressing cells and with 5mM vactosertib. **h**, as in **g**, treated with 5 ng/ml TGF-β1 (**g** and **h**, n = 3). **i**, Albumin proteolysis measured by fluorescence microscopy of self-quenched bovine serum albumin (DQ-BSA) signal in MCF10A ERK2^D319N^ expressing cells and with 5mM vactosertib. **j**, as in **i**, treated with 5 ng/ml TGF-β1 (**i** and **j**, n = 3). **k**, Electron micrographs of MCF10A ERK2^D319N^ expressing cells and MCF10A cells treated with 5 ng/ml TGF-β1. White arrows indicate electron dense lysosomes. **l**, Matrigel invasion assay 3 days after 250 ng/ml doxycycline induction with 50 μM chloroquine (CQ), 100 nM concanamycin A (CMA), or 100 nM bafilomycin A1 (BafA) (n = 3). All values are expressed as mean ± SEM (*p < 0.05, **p < 0.01, ***p < 0.001).

Immunostaining of lysosome associated transmembrane proteins Lamp1 and CD63 revealed significant increase in the endolysosomal compartment in both ERK2^D319N^ and TGF-β induced EMT, which was blocked by a TGFBR1 inhibitor in ERK2^D319N^ expressing cells (Fig. 3e,f). We also visualized the expansion of the acidified late endosome and lysosome pool in live cells using lysotracker (Fig. 3g,h). To determine if increased lysosomal staining in cells undergoing EMT is associated with increased lysosomal degradative capacity we measured proteolysis of a self-quenched lysosomal substrate DQ-BSA. Cells engaged in EMT internalized and digested more BSA than control cells (Fig. 3i,j), which was prevented by TGFBR1 inhibition in ERK2^D319N^ expressing cells and was suppressed by addition of inhibitors of lysosomal acidification, chloroquine (CQ) and concanamycin A (CMA) (Extended Data Fig. 3d). Moreover, expression of ERK2^WT^ or ERK2^D319N^ together with TGF-β stimulation increased DQ-BSA digestion when compared to individual stimulation, highlighting the positive cooperativity between these pathways in driving lysosome activity (Extended Data Fig. 3e,f). We corroborated these findings by electron microscopy and observed EMT cells to have an increase in electron dense lysosomes when compared to control cells (Fig. 3k). The induction of the lysosome compartment coincides with increased expression of proteins associated with phagocytosis, actin reorganization and ECM-receptor interactions as cells become invasive (Extended Data Fig. 3g). To evaluate if increased lysosomal activity contributes to the invasive capacity of EMT cells we measured Matrigel transwell invasion and found a significant decrease in invasive capacity when lysosomes were inhibited for the duration of transwell invasion by CQ or vATPase inhibitors concanamycin A (CMA) and bafilomycin A1 (BafA) (Fig. 3l). These data suggest that as cells engage EMT, the lysosomal compartment and associated proteolytic activity increases in a TGFBR1-dependent manner and the increase in lysosomal activity is associated with enhanced invasive capacity.

### EMT associated lysosomal network induction supports amino acid scavenging

We reasoned that lysosomes are induced as part of the reconfiguration of signaling and metabolism observed during EMT. To determine if EMT driven lysosomal induction accompanies increased autophagic flux we assessed LC3B cleavage and lipidation, markers for the induction of autophagy^30^. We observed nearly double the accumulation of the cleaved and lipidated form of LC3B (LC3BII) in EMT compared to control cells after four-hour treatment with CQ (Fig. 4a). Autophagic flux during EMT was also measured using a tandem mCherry-EGFP-LC3B reporter^31^. Fluorescently labeled LC3B becomes localized to autophagosomes when cleaved and lipidated; however, pH-labile EGFP fluorescence is lost following autophagosome-lysosome fusion, while mCherry fluorescence persists allowing for rate calculation (mCherry/EGFP) of autophagic flux. Cells undergoing EMT exhibited a marked increase in autophagic flux that was dependent on lysosomal activity (Fig. 4b). In addition to autophagy, lysosomes also support degradation of cargo acquired via endocytic pathways, including bulk uptake of extracellular cargo via macropinocytosis^17, 18^. Further supporting this idea, we observe increased vacuolization, albumin uptake and digestion, and albumin engulfment into large PI3P-rich vesicles marked by a FYVE-domain fluorescent reporter in our models of EMT (Fig. 1b, 3e,f and Extended Data Fig. 4a), thus we further assessed the ability of these cells to endocytose 70kDa fluorescent dextran via macropinocytosis. We measured dextran accumulation within the endolysosomal compartment marked by DQ-BSA following a 30-minute incubation with dextran and a 60-minute chase in fresh media (Fig. 4c). Dextran uptake was blocked by a 30-minute pretreatment with the NHE1 inhibitor 5-(N-ethyl-N-isopropyl)-Amiloride (EIPA) (Fig. 4d). Taken together these data show that as cells undergo EMT and increase motility they induce the lysosomal compartment and elevate scavenging capacity.

**Fig. 4.**
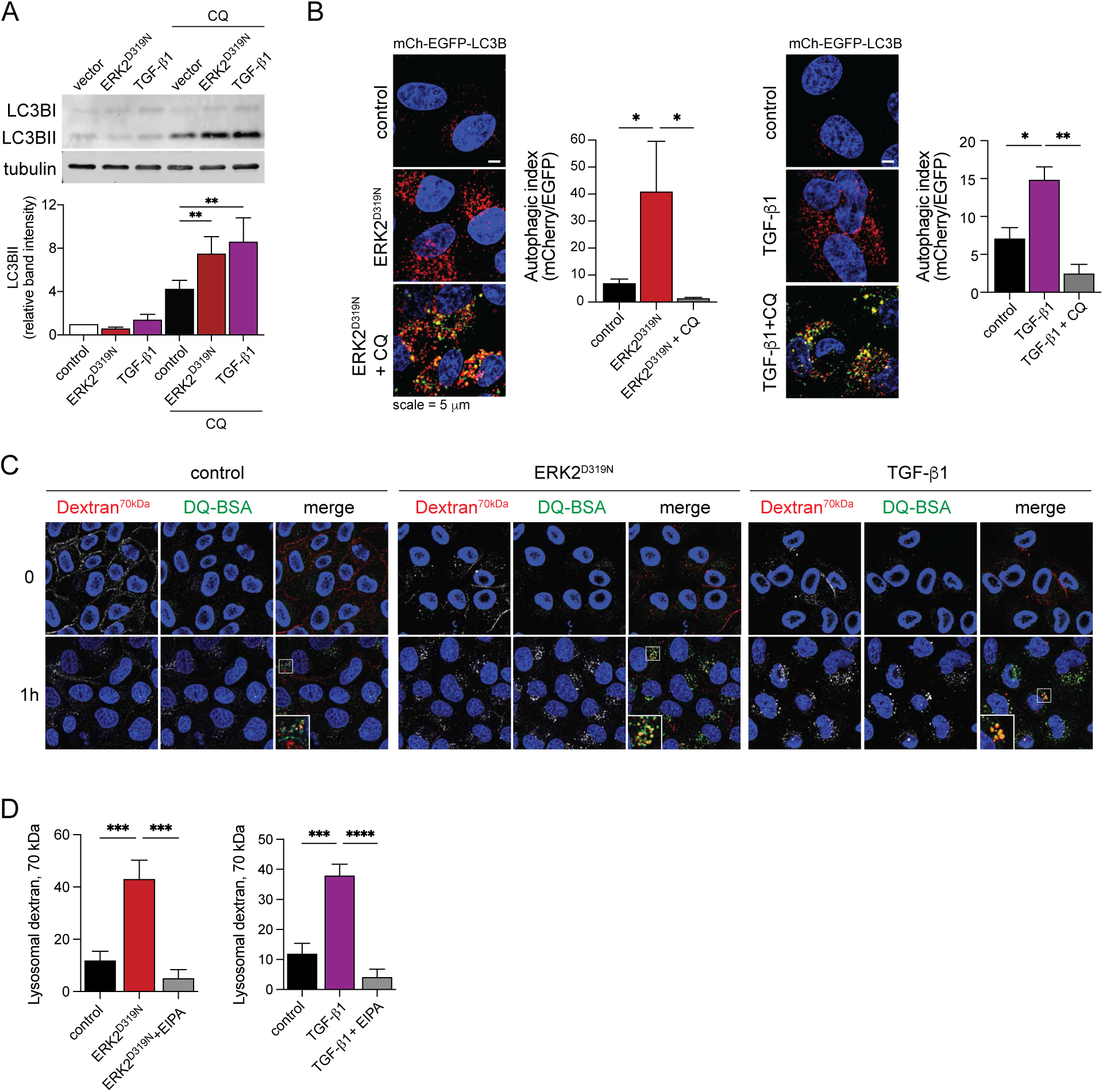
EMT associated lysosomal network induction supports amino acid scavenging. **a**, Immunoblot of MCF10A ERK2^D319N^ expressing cells and TGF-β1 (5 ng/ml) treated cells and after 4-hour treatment with 50μM chloroquine (CQ), mean ± SD (n = 3). **b**, Live cell fluorescence microscopy of mCherry-EGFP-LC3B expressing MCF10A cells expressing ERK2^D319N^ or treated with 5 ng/ml TGF-β1. Also treated with 50 μM chloroquine (CQ) for 4 hours (n = 3). **c**, Live cell fluorescence microscopy of dextran (70kDa) uptake in ERK2^D319N^ expressing or TGF-β1 (5 ng/ml) treated MCF10A cells. Lysosomes were marked by DQ-BSA. Inserts show enlarged area indicated by white box. **d**, Relative dextran uptake corresponding to (C) in control and EMT cells and those pretreated for 30 min with 25μM 5-(N-ethyl-N-isopropyl)-Amiloride (EIPA) (n = 5). Values expressed as mean ± SEM, unless specified (*p < 0.05, **p < 0.01, ***p < 0.001, ****p < 0.0001).

### EMT is marked by a reconfiguration of amino acid transport and increased amino acid storage

We reasoned the metabolic shift and increased scavenging observed during EMT could be coupled to altered amino acid transport, thus we interrogated our EMT proteomics data for changes in transporters primarily located at either the plasma membrane (PM) or endomembranes. The lysosomal glutamine transporter SNAT7 (SLC38A7) required for macropinocytosis driven proliferation in conditions lacking amino acids^32^ was elevated during EMT, while the PM glutamine transporter ASCT2 (SLC1A5) and the essential AA (EAA) heteromeric transporter CD98 comprised of LAT1 and 4F2hc (SLC7A5 and SLC3A2), were all suppressed (Fig, 5a,b). Despite decreased expression of PM associated transporters, the level of free amino acids was elevated 30-50% in EMT cells where protein scavenging is elevated compared to control cells and this increase was reversed by lysosomal inhibition with CQ (Fig. 5c). Notably, lysosome inhibition did not affect free AA levels in control cells which express higher levels of PM associated transporters.

**Fig. 5.**
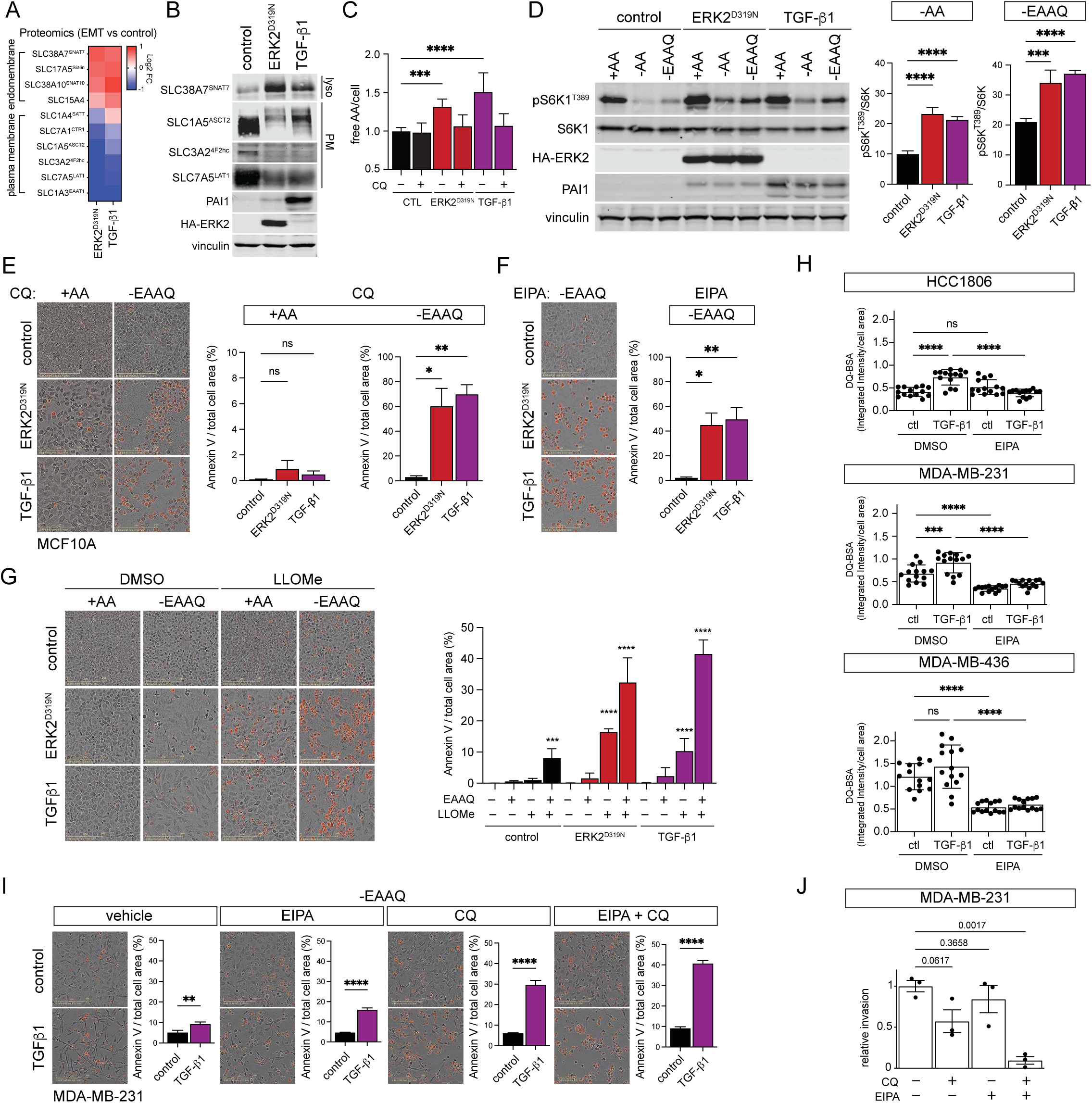
EMT is marked by a reconfiguration of amino acid transport and increased amino acid storage. **a**, Heatmap of log2 fold change in amino acid transporter protein expression in two EMT models compared to control. **b**, Immunoblot of MCF10A amino acid expression in controls cells and EMT. **c**,Quantification of free amino acids per cell in control cells and those undergoing EMT and with 50 μM chloroquine (CQ) treatment for 24h prior to collection, mean ± SD (n = 3). **d**, Immunoblot of MCF10A control cells and those undergoing EMT starved of all amino acids (-AA) or essential AA and glutamine (-EAAQ) for 60 min. Graphs represent quantification of phosphor-S6K (T389) normalized to total S6K as a percentage of amino acid replete (+AA) controls, mean ± SD (n = 4). **e**, Annexin V staining of MCF10A control cells and those undergoing EMT starved of essential AA and glutamine (-EAAQ) for 48 h in the presence of 50 μM chloroquine (CQ) (n = 3). **f**, Annexin V staining of MCF10A control cells and those undergoing EMT starved of essential AA and glutamine (-EAAQ) for 48 h in the presence of 25 μM 5-(N-ethyl-N-isopropyl)-Amiloride (EIPA) (n = 3). **g**, Annexin V staining of MCF10A control cells and those undergoing EMT pulsed with 1 mM L-Leucyl-L-Leucine methyl ester (LLOMe) for 60 min then washed and placed in replete media or starved of essential AA and glutamine (-EAAQ) for 48 h, mean ± SD (n = 7) across two experiments. **h**, Albumin proteolysis measured by fluorescence microscopy of self-quenched bovine serum albumin (DQ-BSA) signal in triple negative breast cancer cell lines stimulated with TGF-β1 (5 ng/ml) for 4 days and with 25μM 5-(N-ethyl-N-isopropyl)-Amiloride (EIPA), mean ± SD (n = 3). **i**, Annexin V staining of MDA-MB-231 control cells and with TGF-β1 (5 ng/ml) for 4 days starved of essential AA and glutamine (-EAAQ) for 48 h in the presence of 50μM chloroquine (CQ), 25 μM 5-(N-ethyl-N-isopropyl)-Amiloride (EIPA) or both, mean ± SD (n = 3). **j**, Matrigel invasion assay of MDA-MB-231 cells treated with 50uM chloroquine (CQ), 25 μM 5-(N-ethyl-N-isopropyl)-Amiloride (EIPA) or both at the time of seeding in the Boyden chamber. Cells were fixed, stained and cell area measured after 16 h (n = 3). Values expressed as mean ± SEM, unless specified (*p < 0.05, **p < 0.01, ***p < 0.001, ****p < 0.0001).

Activation of mTORC1 is dependent on free AAs at the lysosome^33^ and mTORC1 signaling drives increased cell size and invasion during EMT^34^ (Extended Data Fig. 5a). To determine if increased pools of free AA acquired during EMT are sensed by mTORC1, we removed all AA or EAA and glutamine (-EAAQ) from the culture media for 60 min and assessed mTORC1 activity by measuring S6K phosphorylation. AA starvation led to ∼90% inhibition of mTORC1 and EEAQ reduced activity by ∼80%, while cells engaged in EMT were less sensitive to withdrawal of extracellular AA (Fig. 5d). Next, we sought to determine if AA pools acquired via lysosome mediated scavenging during EMT place a dependency on the lysosomal pathway in this cellular context and found that inhibition of lysosomal degradative function or macropinocytosis sensitized EMT cells to EAAQ withdrawal and resulted in apoptotic cell death (Fig. 5e,f and Extended Data Fig. 5b). Furthermore, lysosomal membranes are liable to permeabilization by lysosomotropic agents including L-Leucyl-L-leucine methyl ester (LLOME) leading to depletion of free AA stores^35^. Cells engaged in EMT were particularly sensitized to transient damage of lysosomal membranes by a 30 min treatment with LLOME prior to washout and incubation in replete or AA deficient media (Fig. 5g). These data indicate that elevated scavenging flux observed during EMT increases cellular AA stores and suggest a potential dependency on increased lysosomal function required to support scavenging mediated nutrient stress resistance.

Breast cancer cells utilize macropinocytosis during nutrient deprivation and several cell lines have been found to exhibit elevated steady-state levels of extracellular fluid-phase uptake^36^. Reflecting our observations in non-tumor MCF10A cells, basal breast cancer cell lines (HCC1806, MDA-MB-231, and MDA-MB-436) also increased albumin uptake in response to prolonged TGF-β treatment and this was reversed by inhibition of macropinocytosis with EIPA (Fig. 5h). Similar to MCF10A EMT models, basal breast cancer cells (MDA-MB-231) treated with TGF-β become, and exhibited increased sensitivity to CQ and EIPA during EAAQ starvation compared to untreated cells, while the combination of CQ with EIPA further elevated apoptotic cell death suggesting an increasing dependence on lysosomal function and macropinocytosis (Fig. 5i). The EMT program is associated with increased invasive capacity and basal breast cancer cells (e.g MDA-MB-231) represent an invasive breast cancer model. Addition of CQ or EIPA individually reduced the number MDA-MB-231 or EMT-induced MCF-10A cells able to invade through Matrigel (Fig. 5j), while the combination of the two drugs further suppressed invasion. These data suggest that cells that become invasive through EMT associated signaling may also develop a dependence on scavenged amino acids which are required to fuel cell invasion.

### c-MYC opposes EMT associated amino acid scavenging and limits cell invasion

Cells switch from a proliferative to an invasive program during EMT and this was reflected in the downregulation of Hallmark gene sets associated with the transcription factor and protooncogene c-MYC, a potent driver of cell proliferation (Fig. 1e,f). In agreement, a decrease in c-MYC protein levels and loss of the stabilizing phosphorylation of c-MYC Ser62 was observed in cells undergoing EMT (Fig. 6a), as was a suppression of c-MYC regulated proteins (Fig. 6b). Several lines of evidence highlight the broad importance of c-MYC in the EMT transition as related to our study. First, the TGF-β pathway suppresses c-MYC^37, 38^. Second, c-MYC promotes expression of glycolysis and plasma membrane located AA transporter genes^39–43^, and third, c-MYC suppresses lysosomal gene expression^27, 44^. Together these observations prompted us to test whether c-MYC expression would reverse the EMT associated phenotypes described above. Inducible expression of c-MYC resulted in suppression of the TGF-β response as measured by PAI1 expression, induction of lysosomal markers and reversed the AA transporter switch (Fig. 6c). This was further corroborated by c-MYC mediated suppression of CD63, LAMP2, and the lysosomal glutamine transporter SNAT7, which are all upregulated during EMT (Fig. 6d and Extended Data Fig. 6a,b). Conversely, c-MYC induction restored plasma membrane expression of the Gln transporter ASCT2 (Fig. 6e) and the EAA transporter LAT1 (Extended Data Fig. 6c) in cells undergoing EMT. Similarly, c-MYC expression suppressed lysotracker signal (Fig. 6f) and cathepsin B activity measured by Magic Red assay (Fig. 6g) in EMT cells. As described above, the lysosomal expansion during EMT supports extracellular scavenging. Notably, in addition to suppressing lysosomes, the expression of c-MYC limited the ability of EMT cells to scavenge dextran (Fig. 6h). Moreover, we initially found that ERK2^D319N^ expression dramatically limited the glucose response of MCF10A cells (Fig. 1j), thus we posited that c-MYC suppression during EMT is likely associated with the observed metabolic shift. Indeed, c-MYC positively regulates genes associated with elevated aerobic glycolysis^45^ and the induction of c-MYC in MCF10A cells nearly doubled the glycolysis rate measured by ECAR while MYC expression in ERK2^D319N^ expressing cells restored the glycolysis rate to that of control cells (Fig. 6i and Extended Data Fig. 6d). The inverse was observed when we measured mitochondrial oxygen consumption rate (OCR), where c-MYC suppresses the maximal OCR in control cells and the induced rate of ERK2^D319N^ expressing cells (Fig. 6j and Extended Data Fig. 6e). c-MYC has been shown to limit breast cancer cell invasion and the number of metastases in nude mice^46^ and similarly we find c-MYC expression interfered with the ability of EMT-engaged MCF10A cells to invade through Matrigel (Fig. 6k). Together, these data highlight the coordinated regulation of cell metabolism, nutrient transport and endolysosomal system by EMT associated signaling and c-MYC.

**Fig. 6.**
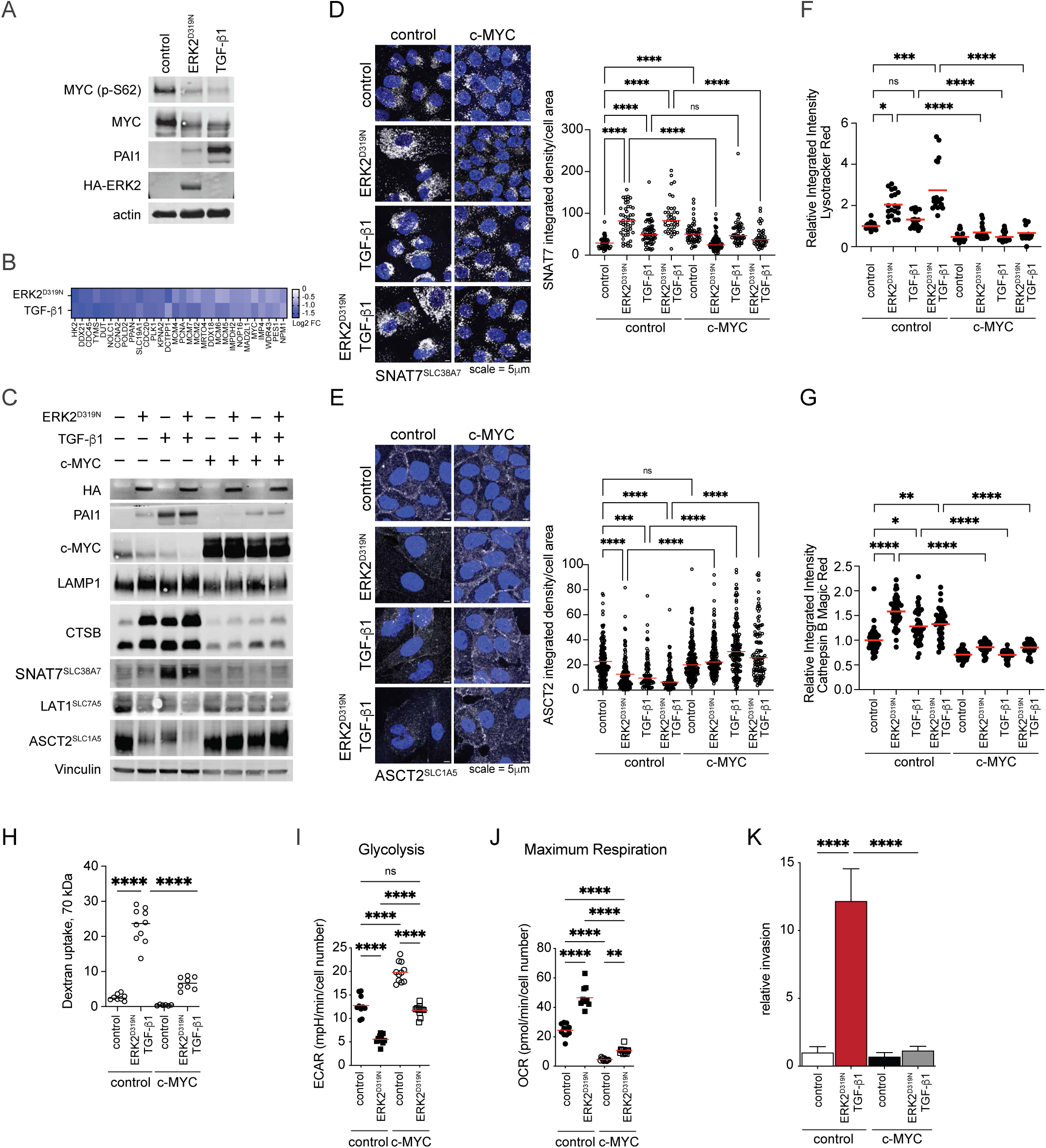
c-MYC opposes EMT associated amino acid scavenging and limits cell invasion. **a**, Immunoblot of MCF10A control cells and those undergoing EMT. **b**, Heatmap of MYC regulated protein expression changes in two EMT models compared to control. **c**, Immunoblot comparing MCF10A undergoing EMT with c-MYC expression. **d**, Immunofluorescence of SNAT7 expression in MCF10A undergoing EMT with c-MYC expression. Data collected over 3-7 fields, n ≥ 38 cells per group. **e**, Immunofluorescence of ASCT2 expression in MCF10A undergoing EMT with c-MYC expression. Data collected over 4-5 fields, n ≥ 23 cells per group. **f**, Relative lysotracker signal intensity in MCF10A undergoing EMT with c-MYC expression (n = 4). **g**, Relative Cathepsin B Magic Red activity in MCF10A undergoing EMT with c-MYC expression (n = 9). **h**, Relative TMR-dextran (70 kDa) uptake in MCF10A undergoing EMT with c-MYC expression, data collected over 8-10 fields, n > 80 cells per group. **i**, Glycolysis rate (ECAR) measured by seahorse analyzer in MCF10A undergoing EMT and with c-MYC expression (n ≥ 11). **j**, Maximum oxygen consumption (OCR) measured by seahorse analyzer in MCF10A undergoing EMT and with c-MYC expression (n ≥ 11). **k**, Matrigel invasion assay of MCF10A cells expressing ERK2^D319N^, c-MYC, or both stimulated by doxycycline (0.5 ng/ml) and treated with TGF-β1 (5 ng/ml) for 2 days prior to seeding in the Boyden chamber. Cells were fixed, stained and cell area measured after 24 h, mean ± SD (n = 6) from two experiments. (*p < 0.05, **p < 0.01, ***p < 0.001, ****p < 0.0001).

## Discussion

Two hallmarks of EMT are increased invasive capacity and improved stress resistance. Our study demonstrates that EMT induction by cooperative ERK2 and TGF-β signaling increases cellular scavenging of extracellular proteins and improves nutrient stress resistance. We find that EMT associated c-MYC suppression initiates a shift away from glycolysis to increases in mitochondrial respiration, macropinocytosis and lysosomal activity. This shift enhances cellular resilience against amino acid deprivation, while also increasing cellular capacity to invade and migrate. Previous studies have shown lysosomes support early EMT events, including loss of cell polarity, through lysosome-mediated degradation of E-cadherin^47–49^, while autophagy-associated lysosomes limit prolonged EMT and suppress apoptosis^50^. During nutrient starvation, cells increase lysosome production and activity to support autophagy-mediated nutrient recycling. Extended periods of nutrient deprivation trigger lysosome and autophagy gene induction by the MiTF/TFE-family of transcription factors^51^. In contrast, we observed EMT-associated lysosome induction occurs even in nutrient-replete conditions, suggesting an ancillary mode of lysosomal regulation essential for cell invasion. Furthermore, the kinases ERK2 and mTORC1 are required for EMT-associated cell invasion and show elevated signaling in our models^9, 34^. Importantly, both ERK2 and mTORC1 have been found to phosphorylate and inhibit the MiTF/TFE-family, thereby blocking lysosome and autophagy gene induction in nutrient replete condtions^51, 52^. The elevated ERK2 and mTORC1 signaling observed in our models suggests a decoupling of their inhibitory effect on TFE-family mediated lysosomal regulation during EMT. Pancreatic ductal adenocarcinoma cells that become reliant on autophagy and macropinocytosis also exhibit elevated mTORC1, while simultaneously maintaining MiTF/TFE activity, enabling anabolic pathway activation and rapid nutrient stress adaptation^53^. Additionally, c-MYC suppresses lysosomal gene expression via direct competition for promoter occupancy of MiTF/TFE target genes^27, 44^. During EMT we observed decreased c-MYC protein expression and decreased expression of c-MYC driven genes suggesting a potential de-repression of lysosomal genes. In support of this idea the induced expression of c-MYC in MCF10A cells undergoing EMT dampened lysosomal abundance and activity.

The high anabolic rate required to fuel growth of RAS and PI3K driven tumors is significantly supported by macropinocytosis, especially in nutrient limited conditions^17, 54, 55^. We find that direct EMT activation leads to elevated rates of macropinocytosis in non-tumor breast cells and resistance to amino acid depravation, while recent studies have found that prolonged glutamine or leucine starvation of macropinocytosis competent pancreatic cancer cells led to activation of EMT and increased invasive capacity^20, 21^. Placing our findings in this context suggests that nutrient scavenging and EMT programs converge to increase cellular fitness in response to amino acid limiting conditions, highlighting an underappreciated resilience mechanism associated with EMT, in addition to those ascribed to chemoresistance and hypoxia adaptation^56, 57^. Furthermore, we find that in addition to increased scavenging MCF10A cells undergoing EMT have increased stores of free amino acids providing a potential mechanism of resilience in these cells, which was disrupted by lysosomal inhibition. Relatively little is known about how cells manage their amino acid stores, however recent work suggests that mammalian cells sense and respond rapidly to loss of extracellular amino acids by increasing lysosomal retention of specific amino acids^35^. While in budding yeast, the luminal pH of the lysosome-like vacuole oscillates as cells transit the cell cycle controlling amino acid storage, protein translation, and cell size^58^. Whether these processes are similarly balanced in mammalian cells, which also integrate the ability to migrate, remains to be elucidated. Notably, EMT programs inhibit or slow the cell cycle; however, we expect that these alterations, along with increased flux through endocytic pathways as cell engage migration machinery and changes in amino acid transport, collectively contribute to the increased amino acid storage observed in our study.

Here we show that cells engaging EMT-signaling reconfigure how they acquire amino acids and enhance resilience against potential periods of nutrient deprivation. Similarly, preimplantation embryos exhibit robust metabolic plasticity marked by their ability to survive and proliferate in the absence of free amino acids and a recent study found that mouse embryonic stem cells rely on macropinocytosis for amino acid acquisition mediated by lysosomal induction and suppression of plasma membrane amino acid transporters^59^. As cells exit pluripotency this balance shifts, lysosomes are suppressed, and plasma membrane transporters are induced. The direct role of c-MYC in reconfiguring stem cell amino acid transport has not been explored, but c-MYC function is likely to be modulated, allowing for lysosomal induction while maintaining pluripotency^45^. When considering a phenotypic shift from proliferation to migration in breast cancer cells, decreasing c-MYC levels are associated with increased integrin expression, invasion and metastasis, while elevated c-MYC levels are associated with rapid proliferation and suppression of integrins^46^. Notably, migrating cells traffic integrins via circular dorsal ruffles and macropinocytosis^60^, a mechanism recently found to be regulated by AXL, an EMT and metastasis associated receptor tyrosine kinase^61^. Furthermore, our model also captures a shift away from high glycolysis to increased mitochondrial respiration, which was recently observed when comparing primary tumor cells to those found in nascent lung metastases in mouse studies of human triple negative breast cancer patient derived xenografts^62^. In addition, increased mitochondrial respiration and biogenesis has been associated with EMT, breast tumor cell invasion, survival in circulation, and distant metastasis^63^. The integration of nutrient acquisition through macropinocytosis with cell migration and nutrient stress survival described here is further supported by reports of altered nutrient signaling with increased invasion^64^ and the ability of starved epithelial cells to scavenge extracellular matrix proteins for survival^65^. Together, our findings establish that cells reconfigure fundamental metabolic architecture when transitioning from a highly proliferative state to an invasive one, and that EMT associated signaling and c-MYC regulation hold a central role in balancing this transition.

Efforts to exploit macropinocytosis to improve drug delivery are ongoing^66^, including mechanistic insights into the delivery of albumin conjugated drugs^67^. Indeed, nanoparticulate albumin bound paclitaxel (nab-PTX, Abraxane) uptake is enhanced by macropinocytosis, which in turn can be stimulated by fasting or drug mediated nutrient starvation mimicry in RAS mutated cancers^68^. Critically, ERK inhibition suppressed macropinocytosis in these cells and blocked drug conjugate uptake and efficacy, highlighting the importance of ERK signaling on cargo uptake by macropinocytosis. Considering our findings the identification of drug conjugates selectively toxic to tumor cells engaged in EMT might further complement strategies most effective against rapidly dividing cells and help eradicate drug tolerant persister cells that cycle between proliferative and dormant or invasive states^22^.

## Methods

### Cell lines and culture conditions

MCF10A cells were a gift from Joan Brugge (Harvard Medical School), MDA-MB-231 (HTB-26), HCC1806 (CRL-2335), and MDA-MB-436 (HTB-130) were purchased from ATCC. HEK293T (Q401) were purchased from GenHunter. MCF10A cells were cultured in DMEM:F12 medium (Corning 10-092-CV) supplemented with 5% horse serum (Gibco 16050-122), 10 μg/ml insulin (Sigma-Aldrich I9278), 100 ng/ml cholera toxin (Sigma-Aldrich C8052), 20 ng/ml epidermal growth factor (PeproTech AF-100-15) and 0.5 mg/ml hydrocortisone (Sigma-Aldrich H0888). Cells were cultured in in high-glucose DMEM (Gibco 11965-118) supplemented with 10% FBS (Sigma-Aldrich F0926), apart from HCC1806 which were cultured in RPMI 1640 (Corning 10-040-CV) with 10% FBS. All cells were maintained in a 5% CO2 humidified incubator and passaged every 2–3 days. All cell lines were routinely tested for mycoplasma contamination using MycoAlert (Lonza LT07-318) during culturing.

During time course experiments cells were passaged every two days, seeded at 30-40% confluence (1 x 10^6^ cells per 10 cm dish), and maintained in 250 ng/ml doxycycline when expressing pTRIPZ-HA-ERK2 (PURO). Cells harvested at 24 h were seeded 48 h prior and treated with doxycycline ∼ 24h after seeding. Experiments utilizing pINDUCER20-HA-ERK2 (NEO) with pLKO.1-puro or TRE-MYC-P2A-EGFP-PURO^69^ were maintained at 500 ng/ml doxycycline.

Amino acid starvation media (DMEM:F12 or high-glucose DMEM lacking all amino acids) was prepared by the Memorial Sloan Kettering Cancer Center Media Core Facility. Horse serum was dialyzed using a 10,000 molecular weight cut-off dialysis flask (Thermo 87762). Media were replenished using dialyzed FBS (Sigma F0392) or dialyzed horse serum, MEM amino acid solution (Thermo 11130051), non-essential amino acid solution (Thermo 11140050) and L-glutamine solution (Sigma 59202C).

### Plasmid construction

PCR for plasmid construction of HA-ERK2 wild-type and corresponding mutants (D319N and Y261A) was performed using Phusion polymerase (NEB) from previously published pBABE-HA-ERK2 templates^9^ and products were ligated with T4 ligase (NEB) following manufacturers protocol into pTRIPZ-tetON-RFP^70^ digested with AgeI and ClaI or pENTR1A no ccDB (Addgene, Cat# 17398), as described by Gomes and collegues^10^ to generate pINDUCER20-HA-ERK2 and pINDUCER20-HA-ERK2^D319N^. Mutagenesis of pENTR1A-HA-ERK2 (E320K and E79K) was performed using QuikChange II XL site directed mutagenesis kit (Agilent) following the manufacturers protocol. The following mutagenesis primer sequences were used; ERK2-E320K, 5’-AGCAGTATTATGACCCAAGTGATAAGCCCATTGCTGAA-3’ and 5’-TTCAGCAATGGGCTTATCACTTGGGTCATAATACTGCT-3’, ERK2-E79K, 5’-CTACTGCGCTTCAGACATAAGAACATCATCGGCA-3’ and 5’-TGCCGATGATGTTCTTATGTCTGAAGCGCAGTAG-3’. pENTR1A-HA-ERK2 (E320K and E79K) and pENTRY-GFP (Addgene, Cat# 15301) were recombined into the Gateway destination vector pInducer20 (Addgene, Cat# 44012) using LR clonase II (Life Technologies). Vectors were verified by sequencing.

### Generation of virus transduced stable cell lines

HEK293T cells were co-transfected with the lentiviral plasmid of interest with pVSVG and delta8.9 using X-tremeGENE HP DNA transfection reagent (Roche). The following viral vectors were used in this study: a series of doxycycline-inducible HA-tagged ERK2 expression vectors, including pTRIPZ-tetON-HA-ERK2 (PURO), pTRIPZ-tetON-HA-ERK2^Y261A^, and pTRIPZ-tetON-HA-ERK2^D319N^ (all generated in this study). The pINDUCER20-based constructs included pINDUCER20-HA-ERK2 (NEO) and pINDUCER20-HA-ERK2^D319N^ (Gomes et al., 2019, ref. 10), as well as pINDUCER20-HA-ERK2^Y261A^, pINDUCER20-HA-ERK2^E79K^, and pINDUCER20-HA-ERK2^E320K^ (this study). The TRE-MYC-P2A-EGFP-PURO plasmid was obtained from Gardner et al. 2024 (ref. 69). Additional expression vectors obtained from Addgene included LPCX-Smad2 (Cat# 12636), LPCX-Smad3 (Cat# 12638), LPCX-Smad2 deltaSSMS (Cat# 12637), and LPCX-Smad3 deltaSSVS (Cat# 12639). Autophagy reporter constructs included pBABE-puro mCherry-EGFP-LC3B (Addgene, Cat# 22418) and pLenti-EGFP-2xFYVE (Addgene, Cat# 136996). To achieve gene knockdown, lentiviral pLKO.1-puro shRNA constructs were obtained from Sigma-Aldrich targeting GFP (shGFP, TRCN0000072181), SMAD2 (shSMAD2 #1, TRCN0000040037; shSMAD2 #2, TRCN0000288592), SMAD3 (shSMAD3 #1, TRCN0000330127; shSMAD3 #2, TRCN0000330128), and SMAD4 (shSMAD4 #1, TRCN0000010321; shSMAD4 #2, TRCN0000010323). Virus-containing supernatants were collected at 48 hr after transfection. Cells were infected with 0.45 mm PVDF membrane-filtered viral supernatant in the presence of 8 mg/ml polybrene for 24 hr and then replaced in fresh media. At about 24-48 hr post-infection, puromycin (2 mg/ml), G418/neomycin (300 mg/ml) was added to select positively transduced cells for 2-5 days.

### Antibodies

The following primary antibodies were used for western blotting (WB) and immunofluorescence (IF) at the indicated dilution: anti-E-cadherin (BD Biosciences, Cat# 610181; WB, 1:1,000), anti-Fibronectin (BD Biosciences, Cat# 610077; WB, 1:1,000), anti-PAI-1 (BD Biosciences, Cat# 612024; WB, 1:1,000), and anti-CD63 (H5C6) (BD Biosciences, Cat# 556019; IF, 1:100). Antibodies from Cell Signaling Technology included anti-phospho-p44/42 MAPK (Erk1/2) (Thr202/Tyr204) (D13.14.4E) (Cat# 4370L; WB, 1:1,000), anti-p44/42 MAPK (L34F12) (Cat# 4696S; WB, 1:1,000), anti-phospho-p70 S6 kinase (Thr389) (Cat# 9205L; WB, 1:1,000), anti-p70 S6 kinase (49D7) (Cat# 2708S; WB; 1:1,000), anti-c-Myc (Cat# 9402S; WB, 1:1,000), anti-phospho-c-Myc (Ser62) (E1J4K) (Cat# 13748S; WB, 1:1,000), anti-phospho-p90RSK (Ser380) (D5D8) (Cat# 12032S; WB, 1:1,000), anti-LAT1, SLC7A5 (E9O4D) (Cat# 32683S, WB; 1:1,000, IF 1:200), anti-ASCT2, SLC1A5 (D7C12) (Cat# 8057S; WB, 1:1,000, IF 1:200), anti-4F2hc, CD98, SLC3A2 (D6O3P) (Cat# 13180S; WB, 1:1,000), anti-phospho-Smad2 (Ser465/467) (138D4) (Cat# 3108S; WB, 1:1,000), anti-phospho-Smad3 (Ser423/425) (Cat# 9523S; WB, 1:1,000), anti-Smad2 (L16D3) (Cat# 3103S; WB, 1:1,000), anti-Smad3 (C67H9) (Cat# 9523S; WB, 1:1,000), anti-Smad4 (Cat# 9515S; WB, 1:1,000), anti-LC3B (D11) (Cat# 3868S; WB, 1:1,000), anti-Cathepsin B, CTSB (D1C7Y) (Cat# 31718S; WB, 1:1,000), and anti-LAMP1 (Cat# 9091S; WB, 1:1,000, IF, 1:100). Antibodies from Santa Cruz Biotechnology included anti-HA-probe (F-7) (Cat# sc-7392; WB, 1:1,000), anti-ZEB1 (H-102) (Cat# sc-25388; WB, 1:500), anti-LAMP2 (Cat# sc-18822; IF, 1:100), anti-Cathepsin C, CTSC (Cat# sc-74590; WB, 1:500), and anti-RSK1 (C-21) (Cat# sc-231G; WB, 1:1,000). Sigma-Aldrich supplied anti-α-Tubulin (Cat# T6199; WB, 1:1,000), anti-Vinculin (Cat# V9264; WB, 1:1,000), and anti-SLC38A7, SNAT7 (Cat# HPA041777; WB, 1:1,000, IF, 1:100). Anti-phospho-Smad3 (Ser423/425) (EP823Y; WB, 1:1,000) was purchased from Abcam (Cat# ab52903).

### Immunoblotting

Cells were washed once with PBS and lysed with 10% TCA lysis solution (10% trichloroacetic acid, 25 mM NH4OAc,1 mM EDTA, 10 mM Tris-HCl, pH 8.0). Precipitated lysates were scraped using a silicon scraper, centrifuged at top speed (5-10 min, 4°C) and supernatant aspirated. Pelleted precipitated proteins were neutralized and resolubilized in a 0.1 M Tris-HCl (pH 11) resuspension solution containing 3% SDS and boiled for 10 min. Protein concentrations of resolubilized lysates were determined by the DC Protein Assay Kit II (Bio-Rad Laboratories), normalized with addition of resuspension solution and 5X Laemmli loading buffer. Samples (20 ug total protein per well) were resolved by SDS–polyacrylamide gel electrophoresis under reducing conditions. Proteins were transferred from the gels to nitrocellulose membranes (GE Healthcare Systems) electrophoretically and then membranes were blocked in Tris-buffered saline-based Odyssey Blocking buffer (LI-COR Biosciences). Membranes were incubated with the primary antibodies overnight at 4 °C in TBS Odyssesy Blocking buffer, washed 3x in Tris-Buffered Saline with Tween-20 (TBST) and incubated with IRDye 800CW and 680RD conjugated secondary antibodies (LI-COR Biosciences) diluted 1:10,000 in Odyssey Blocking buffer for 1h at room temperature. All incubation steps and washes were performed with gentle rocking. Immunoblots were imaged using an Odyssesy CLx scanner (LI-COR Biosciences).

### Gene Expression Analysis

Vector control, D319N ERK2, and Y261A ERK2 expression was induced by doxycycline in Tet-on MCF10A cell systems. RNAs were purified from three biological replicates of cells for each condition that were grown for 1, 3, 5, 7 and 9 days using an RNA isolation kit (Qiagen). Partners Healthcare Personalized Medicine performed a microarray using GeneChip Human Transcriptome Array 2.0 (Affymetrix). The CEL files were read into R (R Core Team (2020). R: A language and environment for statistical computing. R Foundation for Statistical Computing, Vienna, Austria. URL https://www.R-project.org/) with the help of the oligo package^71^, the Platform Design Info for Affymetrix HTA-2_0 package (pd.hta.2.0; http://www.bioconductor.org/packages/release/data/annotation/html/pd.hta.2.0.html), and the annotation package hta20transcriptcluster.db (http://www.bioconductor.org/packages/release/data/annotation/html/hta20transcriptcluster.db.html).

The raw intensity values were normalized with the Robust Multichip Average algorithm^72^ as implemented in the oligo::rma function. Statistically significantly differentially expressed genes were determined with limma^73^, comparing the values of the ERK2^D319N^ (DN) replicates to those of the control conditions (Empty Vector - EV, ERK2^Y261A^ - YA) for every day of the time course, using an adjusted p-value threshold of 5%. The resulting log2-transformed fold change (logFC) values were used for downstream analyses and visualizations. In addition, we applied the Transcript Time Course Analysis algorithm as implemented in the TTCA package to identify genes that change when taking the entire time course into consideration^74^. The microarray data were deposited in the Gene Expression Omnibus (GEO) database with accession no. GSEXXXXXX-pending.

### Proteomics Analysis

#### Sample processing for proteomic analysis

Four biological replicates were collected for the time course of ERK2-driven EMT (Fig. 1d,f). Two sets of replicates were lysed using a detergent based lysis method and two sets were prepared using TCA precipitation^75^. Three biological replicates were collected when comparing MCF10A control, ERK2^D319N^, and TGF-β1 treated (5ng/ml) cells (Extended Data Fig. 3a,b) and protein samples were extracted using TCA precipitation.

#### Detergent protein extraction

Cells were homogenized for 30 s in 5 ml of lysis buffer (2% SDS, 150 mM NaCl, 50 mM Tris-HCl pH 8.5, Roche mini-complete EDTA free protease inhibitors, 5 mM DTT) in 50 ml conical tubes using a Biospec Tissue Tearor homogenizer. Cell homogenates were incubated at 60° C for 45 min, allowed to return to room temperature, alkylated with 14 mM (final concentration) iodoacetamide for 45 min in the dark, then quenched with an additional 5 mM DTT. Protein extracts were methanol/chloroform precipitated, transferred to 2 ml tubes, and washed three times in ice-cold acetone. The protein pellets were resolubilized in 1.5 ml 8 M urea buffered with 50 mM Tris at pH 8.5 using the homogenizer. Insoluble material was pelleted at 15k x g for 10 min and the supernatants were transferred to 50 ml tubes. Protein concentrations were assayed by BCA (Pierce).

#### TCA protein extraction

Cells were pelleted in 10% TCA as previously described^75^ and incubated on ice for 30 min. Cell pellets were washed twice with 10 ml ice-cold acetone. The protein pellets were resolubilized in 400 μl 8 M urea buffered with 10 mM Tris at pH 8.5, 50 μM ammonium bicarbonate and 5 mM EDTA using the homogenizer. Protein concentrations were assayed by BCA (Pierce). For each replicate 3 mg of protein was reduced by 5 mM TCEP for 45 min, alkylated by 12 mM iodoacetamide for 1 h.

Common to both extraction methods above, the urea concentration was lowered to 2 M by adding three volumes of digestion buffer (10mM Tris pH 8.5, 1mM CaCl2). Protein extracts were digested by adding lysyl endopeptidase (lysC, Wako Chemical) at 1:100 ratio (enzyme:substrate) and incubated overnight at 30° C. They were diluted to 1 M urea with one volume digestion buffer and digested with sequencing grade Trypsin (Promega) at 1:100 for 6 hours at 37° C. The digestion reactions were quenched by adding neat formic acid to a final concentration of 2%.

Peptides were desalted on Sep-pak C18 solid-phase extraction cartridges (Waters). Prior to drying, a volume corresponding to 100 ug of starting protein was removed to fresh tubes for global protein abundance profiling. All samples were dried overnight in a speed-vac. The 100 ug aliquots of peptides were labeled with 11-plex TMT tags for the ERK2D319N time course experiment or 9-plex for ERK2D319N and TGF-β comparison experiment, following the manufacturer’s protocol for 1 hour and quenched with hydroxylamine. Five ul of each reaction was removed, pooled, diluted to <10% organic solvent, desalted on C18 STAGE Tips, and analyzed with a short run to check labeling efficiency and equal loading.

#### Peptide Fractionation

TMT labeled samples were combined using the tester run results to mix equal total amounts of material from each sample and de-salted on C18 Sep-Pak cartridges. After elution and drying, the TMT mixtures were resuspended in 300 μl buffer A (5% ACN, 10 mM NH4HCO3, pH 8) and separated by high-pH reverse-phase HPLC (56) on a C18 column (Waters, #186003570, 4.6 mm x 250 mm, 3.5 μ ID) using a 50 min gradient from 18% to 38% buffer B (90% ACN, 10 mM NH4HCO3, pH 8) with a flow rate = 0.8 ml/min. Fractions were collected over 45 min at 28 sec intervals beginning 5 min after the start of gradient in a 96-well plate. The original fractions were pooled into 12 samples each containing four fractions (only 48 of 96 fractions were used) by sampling at equal intervals across the gradient, i.e. by combining fractions 1/25/49/73, 3/27/51/75, 5/29/53/77, ….13,37,61,95. The pooled samples were dried down, resuspended in 50 μl of 5% FA, and desalted on STAGE tips. For the TGF-β treatment experiment, 96 fractions were pooled similarly to those described above, but taking 24 pools of four original fractions by combining 1/25/49/73; 2/26/50/74; 3/27/51/75 …. 24/48/72/96.

#### LC-MS/MS Analysis

Samples were re-suspended in 5% formic acid (FA) and analyzed on a Thermo Orbitrap Fusion mass spectrometer (Thermo Fisher Scientific) equipped with an Easy nLC-1000 UHPLC (Thermo Fisher Scientific). Peptides were separated with a gradient of 5–25% ACN in 0.1% FA over 165 min and introduced into the mass spectrometer by nano-electrospray as they eluted off a self-packed 40 cm, 75 μm (ID) reverse-phase column packed with 1.8 μm, C18 resin (Sepax Technologies, Newark, DE). They were detected using the real-time search (RTS) MS3 method^76^ which employs a data-dependent Top10-MS2 method and a RTS triggered SPS-MS3 to collect reporter ions. For each cycle, one full MS scan was acquired in the Orbitrap at a resolution of 120,000 with automatic gain control (AGC) target of 5e5 and maximum ion accumulation time of 100 ms. Each full scan was followed by the individual selection of the most intense ions, using a 3 sec cycle time for collision-induced dissociation (CID) and MS2 analysis in the linear ion trap for peptide identification using an AGC target of 1.5e4 and a maximum ion accumulation time of 150 ms. Ions selected for MS2 analysis were excluded from reanalysis for 60 s. Ions with +1, >+5, or unassigned charge were excluded from selection. Collected MS2 spectra were searched in real time against a database of all uniprot reviewed human sequences (uniprot.org, downloaded May 1, 2019) using binomial score cutoff of 0.75. Passing spectra triggered synchronous precursor selection of up to 10 MS2 ions for HCD fragmentation (CE = 55%) and MS3 reporter ion scans in the orbitrap at resolution = 60,000 using a maximum ion time of 150 ms. The gene close-out feature was enabled and applied to each set of 12 fractions and used to limit the selection and collection of MS3 reporter ions to a maximum of 10 peptides per protein across all 12 runs (ERK_DN) or 24 runs (TGF-β) in each set.

#### Peptide Searching and Filtering and Protein Quantification

MS/MS spectra were searched to match peptide sequences using SEQUEST v.28^77, 78^ and a composite database containing all reviewed human uniprot protein sequences (uniprot.org, downloaded May 1, 2019) and their reversed complement. Search parameters allowed for two missed cleavages, a mass tolerance of 20 ppm, a static modification of 57.02146 Da (carbamidomethylation) on cysteine, and dynamic modifications of 15.99491 Da (oxidation) on methionine and 229.16293 Da on peptide amino termini and lysines. Peptide spectral matches were filtered to a 2% false-positive rate using the target-decoy strategy^79^ combined with linear discriminant analysis (LDA) ^80^ using the SEQUEST Xcorr and ΔCn scores, mass error, charge, and the number of missed cleavages. Further filtering based on the quality of quantitative measurements (reporter ion sum signal-to-noise ≥ 200, isolation specificity >0.7) resulted in a final protein FDR < 1% for all 10plex experiments. Total protein intensity values for each sample were derived from the summed reporter ion intensities from the corresponding channel from all peptides mapping to that protein. Values were normalized to account for small variations in sample mixing based on the sum intensity of values from all proteins in each channel within each 11plex. Relative abundances for each protein across samples within each 11plex were calculated as the fraction of the total intensity derived from each channel.

#### Identifying significantly changing proteins

The protein abundances were loaded into R and re-normalized, using a method established in the Krumsiek Lab. In brief, for every protein, its median abundances across all samples is determined, followed by a dilution factor, which is based on the median-normalized values. For details, see SuppFigS3_px_plots_proteome.Rmd in the github repo.

To identify proteins with statistically significantly changing values across the different conditions, the limma package was used^73^. Missing values were replaced with a pseudo-value (log2(2^min(<observed values>))/2) before limma::lmFit was used to fit a linear model to the observed protein abundances over all replicates and samples of interest. LogFC values were obtained via estimating the coefficients of the model and statistical significantly changing proteins were identified via limma::eBayes() function and limma::topTable() with Benjamini-Hochberg correction for multiple testing and a defined p value cut-off of 0.1 for results presented in Extended Data Fig. 3 and 0.05 for the time course analyses.

#### Clustering

To identify groups of proteins with similar changes over time, we applied kNN-based clustering to the logFC values for DN-vs-YA as calculated by limma (described in previous paragraph): igraph::cluster_walktrap(scran::buildSNNGraph(t(prots.lf), k =40)). Enrichr^81^ curated by the Ma’ayan laboratory was used to generate enrichment plots of KEGG terms from proteomics data.

### Metabolite Profiling and Analysis

Cells were washed once with ice-cold PBS on ice and intracellular metabolites were extracted using 80% (v/v) aqueous methanol. Targeted liquid chromatography-tandem mass spectrometry (LC–MS/MS) was performed using a 5500 QTRAP triple quadrupole mass spectrometer (AB/SCIEX) coupled to a Prominence UFLC HPLC system (Shimadzu) with Amide HILIC chromatography (Waters). Data were acquired in selected reaction monitoring (SRM) mode using positive/negative ion polarity switching for steady-state polar profiling of greater than 260 molecules. Peak areas from the total ion current for each metabolite SRM transition were integrated using MultiQuant v2.0 software (AB/SCIEX). Metabolite raw values were median-normalized (per run) and quotient-normalized (across all samples; according to Dieterle and collegues^82^, missing values were imputed using a kNN-based approach^83^. Limma^73^ was applied to the log2-transformed, normalized values to determine significantly changing metabolites in pairwise comparisons for each day of the time course. Further enrichment analysis of the normalized data was carried out using MetaboAnalyst v4.0, an open-source online platform for comprehensive metabolomics data analysis (https://www.metaboanalyst.ca/).

### Metabolic and bioenergetic assays

For Seahorse XFe96 extracellular flux analysis, 1.0 x 10^4^ vector control and 1.5 x 10^4^ ERK2^D319N^ expressing MCF10A cells were seeded in XF96 cell culture microplates (Agilent, 101085-004) in MCF10A media and incubated in a 5% CO2 incubator at 37°C overnight.

#### Seahorse XF Glycolysis Stress Test

The day after seeding cells, media was removed and wells were washed twice with serum, glucose and pyruvate free RPMI assay media (pH 7.4) (Agilent, 103680-100) supplemented with 2 mM L-glutamine (Sigma Aldrich, G7513), and incubated with 175μL assay media at 37°C for 1 hour in a non-CO2 incubator. During this incubation, Ports A-D of a Seahorse XFe96 FluxPak (Agilent, 103793-100) were loaded with 25 μL of the following compounds dissolved in assay media: Port A – Glucose (Sigma Aldrich, G7021); Port C – Oligomycin A (Sigma Aldrich, 75351); Port D – 2-Deoxy-D-glucose (Sigma Aldrich, D8375). Final concentrations of each compound following injection were: 0.1% DMSO, 10 mM glucose, 2 μM oligomycin A and 100mM 2-Deoxy-D-glucose. Extracellular acidification rate (ECAR) was measured using the Seahorse XFe96 Analyzer. Glycolysis rate was calculated as follows: (maximal rate measurement before oligomycin injection) – (maximal rate measurement before glucose injection).

#### Seahorse XF Cell Mito Stress Test

The day after seeding cells, media was removed and wells were washed twice with serum, glucose and pyruvate free RPMI assay media (pH 7.4) (Agilent, 103680-100) supplemented with 2 mM L-glutamine (Sigma Aldrich, G7513), 1mM sodium pyruvate (Gibco, 11360-070) and 10mM glucose (Sigma Aldrich, G7021), and incubated with 175μL assay media at 37°C for 1 hour in a non-CO2 incubator. During this incubation, Ports A-D of a Seahorse XFe96 FluxPak (Agilent, 103793-100) were loaded with 25 μL of the following compounds dissolved in assay media: Port A – Oligomycin A (Sigma Aldrich, 75351), Port C – carbonyl cyanide-p-trifluoromethoxyphenylhydrazone (FCCP) (Sigma Aldrich, C2920); Port D – rotenone (Sigma Aldrich, R8875) and antimycin A (Sigma Aldrich, A8674). Final concentrations of each compound following injection were: 0.1% DMSO, 2 μM oligoycmin A, 0.5 μM FCCP, and 0.5μM rotenone and antimycin A. Oxygen consumption rate (OCR) was measured using the Seahorse XFe96 Analyzer. Maximal respiration rate was calculated as follows: (final rate measurement after FCCP injection) – (final rate measurement after oligomycin injection).

### Matrigel Invasion assays

MCF-10A cells were trypsinized, collected, and resuspended with resuspension media (DMEM/F12 (Corning), 20% Horse Serum (Gibco)). Resuspension media were aspirated, and cells were resuspended in assay medium (DMEM/F12 (Corning), 0.5% Horse Serum (Gibco), 500 ng/ml hydrocortisone (Sigma), 100 ng/ml cholera toxin (Sigma)). BD BioCoat invasion chambers (Corning) coated with growth factor-reduced Matrigel were used. Invasion chambers were prepared according to the manufacturer’s specifications. Assay medium supplemented with 20 ng/ml EGF (PeproTech AF-100-15) was added to the bottom chamber of the cell culture inserts to serve as the chemoattractant and cells (5 × 10^4^ cells per 250 μl assay medium) were then added to the top chamber and allowed to invade for 24 h. Cells that had migrated to the lower surface of the membrane were fixed with ethanol and stained with 0.2% crystal violet in 2% ethanol.

Transwell invasion assays for MDA-MB-231 cells were performed as described above with minor changes. High-glucose DMEM (Gibco) supplemented with 250 μg/ml BSA (Sigma-Aldrich) was used as the assay medium, and high-glucose DMEM supplemented with 10% FBS (Sigma-Aldrich) and 5ng/ml EGF was used as the chemoattractant (20 h).

Plates were scanned using a digital scanner (Epson) and quantifications were carried out using Fiji/ImageJ2 v2.16^84, 85^. In brief, binary images of the area covered by crystal violet-positive cells was generated using background subtraction, thresholding, and settings that were appropriate for control samples, and these settings were used throughout the analysis. The percentage area covered by crystal violet-positive cells was quantified for each condition, using a minimum of three technical replicates.

### Immunofluorescence

Cells were seeded at a density of 3 x 10^4^ on cover slips (Fisherbrand 12-545-80) in 24 wells. Cells were fixed with 4% paraformaldehyde in PBS for 15 min at room temperature, then rinsed three times with PBS and permeabilized for 10 min in PBS with 0.2% Triton X-100 at room temperature, unless indicated otherwise. Samples were blocked for 1 h using Odyssey TBS Blocking Buffer (LI-COR Biosciences 927-60050), then incubated with primary antibodies overnight at 4° C. The following day, cover slips were washed 3-4 times with PBS and incubated for 1 h with AlexaFluor-488, AlexaFluor-555 or AlexaFluor-647 conjugated antibodies (Thermo Scientific) in 1:2000 dilutions. After washing two times coverslips were incubated for 10 min at room temperature in 1:10,000 DAPI (Biotium) diluted with PBS, then washed 2-3 more times with PBS and mounted using Dako Fluorescence Mounting Medium (Agilent Technologies) onto glass slides (Fisher Scientific). Samples probed for SNAT7 were permeabilized with 0.1% saponin (Sigma-Aldrich SAE0073) for 10 min at room temperature. Target signal intensity was determined by background subtraction, thresholding, and quantified within the boundary area and normalized by cell number, determined by counting nuclei or measured within individual cell boundary using Fiji/ImageJ2 v2.16.

### Live cell imaging

Lysotracker and DQ-BSA analysis was performed on 3 x 10^4^ cells were seeded in 8-well chambered cover slides (Cellvis C8-1.5H-N). The next day 10 μg/ml of DQ-BSA Red (Invitrogen D12051) was added and cells incubated for 18h. 30 min prior to imaging 50 nM Lysotracker Green DND (Molecular Probes L7526) was added, and cells were imaged using Zeiss LSM 800 Laser Scanning Confocal Microscope with Plan Apochromat 40X dry or 63X oil immersion objectives. Higher throughput analysis was conducted on 1.0 x 10^4^ cells seeded in a 96-well plate (Corning) and imaged using Incucyte S3 with the 20x objective.

### Electron microscopy

Samples were washed with PBS then fixed with a modified Karmovsky’s fix of 2.5% glutaraldehyde, 4% paraformaldehyde and 0.02% picric acid in 0.1 M sodium cacodylate buffer at pH 7.2^86^. Following a secondary fixation in 1% osmium tetroxide, 1.5% potassium ferricyanide^87^, samples were dehydrated through a graded ethanol series and embedded in situ in LX-112 resin (Ladd Research Industries). En face ultra-thin sections were cut using a Diatome diamond knife (Diatome, USA, Hatfield, PA) on a Leica Ultracut S ultramicrotome (Leica, Vienna, Austria). Sections were collected on copper grids and further contrasted with lead citrate^88^ and viewed on a JEM 1400 electron microscope (JEOL, USA, Inc., Peabody, MA) operated at 100 kV. Digital images were captured with a Veleta 2K digital camera using RADIUS software (EMSIS, Germany).

### Dextran uptake assay

Cells were seeded at a density of 3 x 10^4^ in 8-well chambered cover slides (Cellvis C8-1.5H-N) and maintained in replete MCF10A media and 70kDa TMR-Dextan (Invitrogen D1818). To quantify uptake and retention of 70kDa dextran and minimize background associated with growth in full media cells were incubated with 1 mg/ml 70kDa dextran and 50 μg/ml DQ-BSA Green (Invitrogen D12050) for 30 min washed five times with PBS and switched to full media and incubated for 60 min prior to live imaging or fixation with 4% paraformaldehyde in PBS for 15 min at room temperature. 50 nM Lysotracker Green DND was added 30 min prior to imaging when used in lieu of DQ-BSA Green. 25 μM EIPA treatments were performed 30 min prior to dextran incubation and maintained during subsequent media changes. Dextran signal intensity was quantified within the boundary area defined by DQ-BSA or lysotracker signal using Fiji/ImageJ2 v2.16.

### L-amino acid assay

Total amount of free L-amino acids (except for glycine) were measured by L-Amino Acid Assay Kit (Cell Biolabs) following the manufacturer’s protocol. Control, ERK2^D319N^ (3 days), and TGF-β1 (5 ng/ml 4 days) MCF10A cells were seeded two days prior to collection on 6 cm dishes (3.5 x 10^3^ cells) and treated with or without 50 μM chloroquine for 24 h. Cells were trypsinized and counted, then 1 x 10^6^ cells resuspended in 1 ml of 1X Assay Buffer and lysed using probe sonication (10 pulses, 30% power, 50% duty cycle) on ice and then centrifuged to remove cell debris. 50 mL of each L-alanine standard or cell lysis was added into wells of a 96-well microtiter plate. Then 50 mL of Reaction Mix was added to each well. The well contents were mixed thoroughly and incubated for 90 minutes at 37° C protected from light. The plate was read with a spectrophotometric microplate reader in the 540-570 nm range. The concentration of L-amino acids was calculated within samples by comparing the sample OD to the standard curve.

### mTORC1 activity assay

ERK2 ^D319N^ expressing or TGF-β1 (5 ng/ml) treated MCF10A cells were seeded (1 x 10^6^) in 10 cm plates two days prior to collection. Cells were washed once with PBS and media exchanged to amino acid replete MCF10A media (+AA), media lacking all amino acids (-AA) or media lacking essential amino acids and L-glutamine (-EEAQ) for 60 min, then collected for immunoblotting. Growth factors and serum were maintained throughout.

### Annexin V assay

ERK2 ^D319N^ expressing or TGF-β1 (5 ng/ml) treated MCF10A or MDA-MB-231 cells were seeded (5.0 x 10^3^ per well) in a 96-well plate (Corning) and grown for 48 h until media change. Cells were washed once with PBS and media exchanged to amino acid replete media (+AA) or media lacking essential amino acids and L-glutamine (-EEAQ), both containing 250 ng/ml annexin V (Biotium) along with either DMSO, 50 μM CQ, 25 μM EIPA, or 100 nM CMA. Alternatively, cells grown for 48 h in a 96-well plate as described above were treated with 1 mM L-Leucyl-L-Leucine methyl ester (LLOMe) for 60 min to permeate lysosomes and release AA stores, then washed three times with PBS and media exchanged to +AA or -EAAQ containing annexin V. In all instances cells were cultured for an additional 48 h and imaged using the Incucyte S3 with a 20x objective. Death was assessed by dividing the area of annexin V signal by total cell area as determined by phase contrast. Growth factors and serum were maintained throughout.

### Magic Red Cathepsin B activity assay

Cathepsin B activity was assessed using the Magic Red assay after three days of EMT induced by 500 ng/ml doxycycline in ERK2 ^D319N^ expressing or/and TGF-β1 (5 ng/ml) treated MCF10A also carrying Tet-inducible c-Myc. Cells were seeded (5.0 x 10^3^ per well) in a 96-well plate (Corning) and grown for 48 h prior to reagent addition. Reagent was added following the manufacturers and imaged after a 30 min incubation using the Incucyte S3 with the 20x objective.

### Statistics and reproducibility

All data are represented as mean ± SEM or mean ± SD, unless otherwise indicated. Statistical significance between groups was determined by unpaired Student’s t-test, Mann-Whitney U-test, and one-way analysis of variance (ANOVA) plus Tukey or Sidak’s multiple comparisons test. Specific statistical test details, including the sample size and significance definition can be found in the legend of each figure. Value of n indicates the number of independent experiments unless otherwise stated. Statistical analyses were performed using GraphPad Prism software (v10.4.0) and R.

## Reporting summary

Further information on research design is available in the Nature Portfolio Reporting Summary linked to this article.

## Data availability

Transcriptomics data have been deposited at GEO: accession number and are publicly available as of the date of publication. Proteomics data have been deposited at MassIVE: accession number and are publicly available. Source data are provided with this paper. All other data supporting the findings of this study are available from the corresponding author on reasonable request.

## Code availability

The code underlying transcriptomics, proteomics and metabolomics analysis is documented within an accompanying R package deposited on https://github.com/abcwcm/Nagiec2025

## Acknowledgements

We thank the members of the Blenis laboratory for critical discussions; the Weill Cornell Medicine Microscopy & Image Analysis Core for confocal microscopy support provided by Janet Sun, Sushmita Mukherjee and Lee Cohen-Gould and electron microscopy & histology services performed by Juan Pablo Jimenez and Lee Cohen-Gould, utilizing the National Institute of Health (NIH) Shared Instrumentation Grant (S10RR027699). The mass spectrometry-based metabolomics work was partially funded by NIH grant 5P01CA120964 (J.M.A.). M.N received NIH T32 training grant support from the National Cancer Institute (NCI) CA009361 and F32 fellowship from the National Institute for General Medical Sciences (NIGMS); GM106582. This work was supported by a grant from NIH/NCI; CA046595 to J.B.

## Author contributions

Conceptualization, M.J.N. and J.B.; Methodology, M.J.N., N.D., F.D., P.M.C., N.K., A.P.M., J.B.N., S.P.G., J.M.A. and J.B.; Investigation, M.J.N., N.D., P.M.C., D.M.S., N.K, J.B.N., S.S., S.O.Y., J.H., A.C., O.M.K., R.K.; Formal analysis, M.J.N., F.D., P.Z., P.M.C, N.D.; Writing—original draft, M.J.N.; Writing—review & editing, M.J.N., A.P.M., B.P., E.E.G., N.D., F.D., and J.B.; Funding acquisition, M.J.N., N.D. and J.B.; Resources, M.J.N., E.E.G., J.M.A., S.P.G., D.B., N.D., and J.B.; Supervision, M.J.N., J.M.A., S.P.G., D.B., N.D., and J.B.

## Extended Data Figure Legends

**Extended Data Fig. 1.**
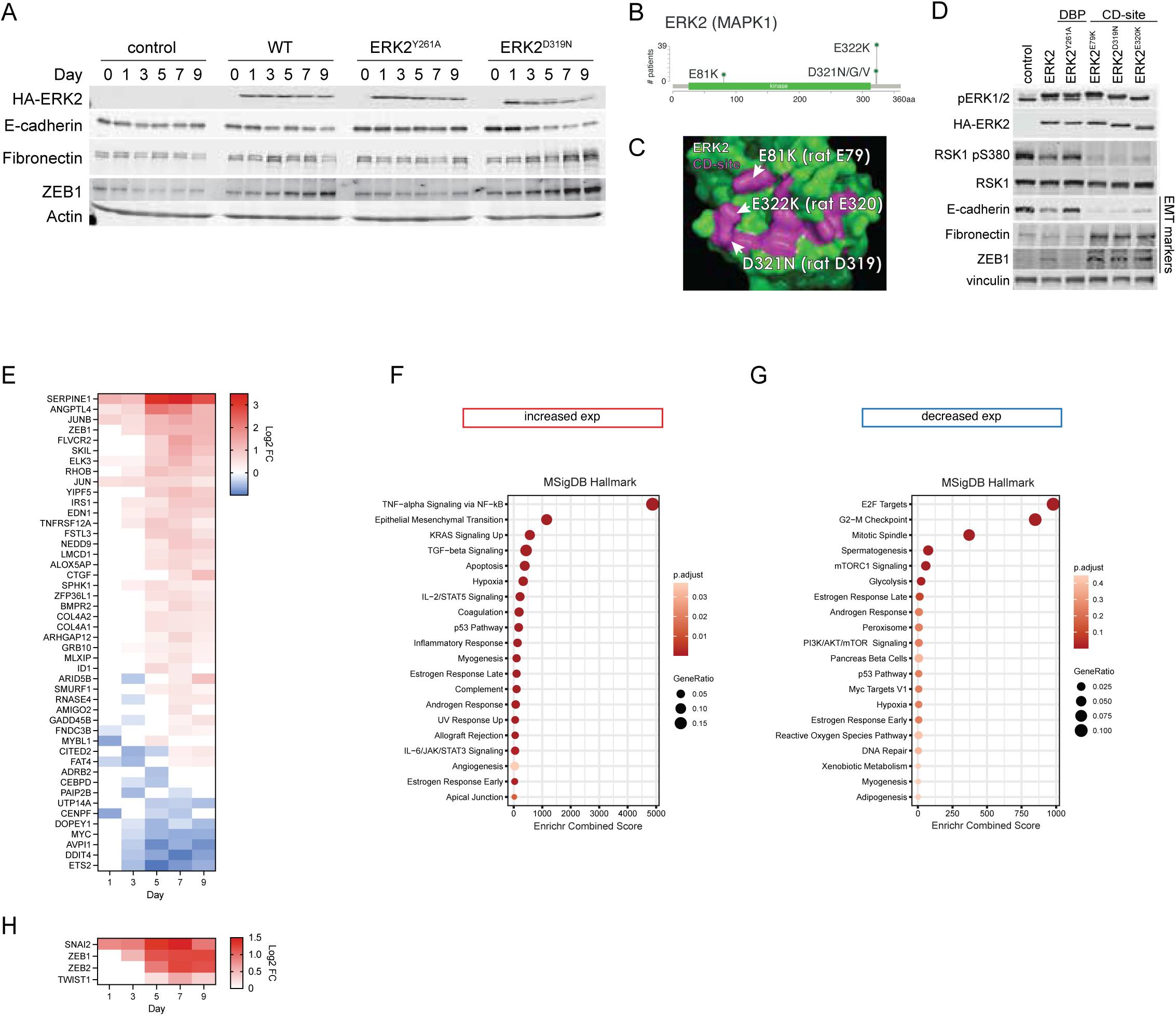
Mutant ERK2 signaling is associated with EMT signatures found in metastasis models. **a**, Immunoblot of MCF10A vector control cells and those expressing ERK2-WT, Y261A, and D319N collected over 9 days of doxycycline stimulation (0.25 ng/ml). Corresponding to images in Fig. 1b. **b**, Mutation diagram of ERK2/MAPK1 common docking site mutation occurrence of non-redundant pan-cancer data available on cBioPortal (101,480 patients). **c**, Three-dimensional rendering of the common docking site of ERK2. Indicated residues correspond to Human numbering along with Rat (constructs used in this study) and are indicated by arrows and colored magenta. **d**, Immunoblot of MCF10A vector control cells and those expressing ERK2 common docking (CD) site mutants found in tumors. **e**, Temporal heatmap of gene expression in MCF10A ERK2^D319N^ expressing cells compared to controls. The gene set was previously identified by Padua and collogues (ref. 24), as the TGF-β response associated with breast tumor lung metastasis. Data represent log2 fold-changes. **f**, Enrichment analysis of transcripts that increase in expression and overlap with Jin and collogues (ref. 25). **g**, Enrichment analysis of transcripts that decrease in expression and overlap with Jin and collogues (ref. 25). **h**, Temporal heatmap of EMT transcription factor gene expression in MCF10A ERK2^D319N^ expressing cells compared to controls.

**Extended Data Fig. 2.**
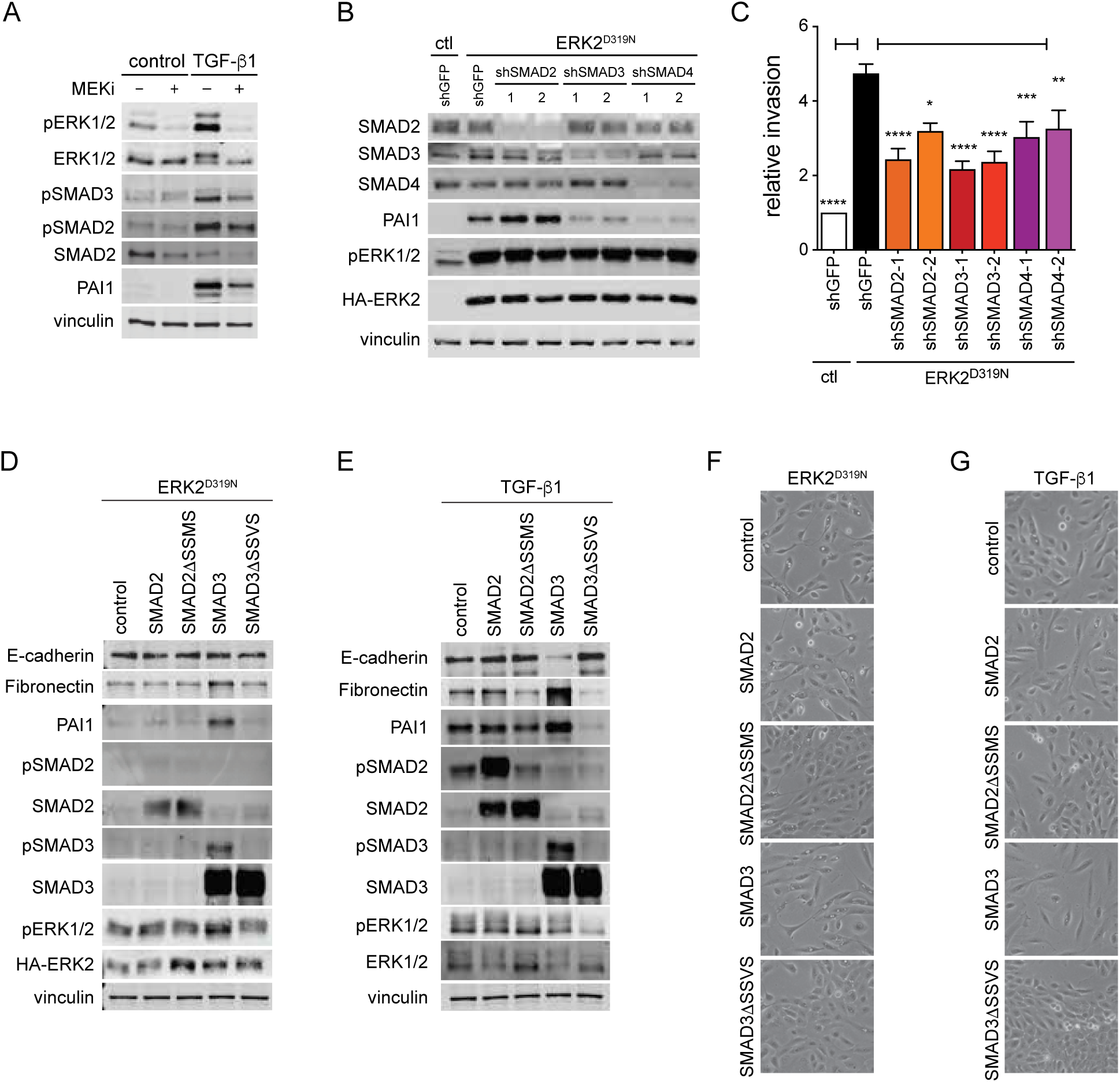
Convergence of ERK2 and TGF-β signaling in EMT. **a**, Immunoblot of MCF10A TGF-β1 (5 ng/ml) stimulated cells (Day 4), treated with 1 μM MEK inhibitor Selumetinib (MEKi) for 24h. **b**, Immunoblot of MCF10A with knockdown of SMAD2, SMAD3, or SMAD4, and effect on PAI-1 expression in MCF10A ERK2^D319N^ expressing cells (Day 3). **c**, Matrigel invasion assay corresponding to samples in **b** (n = 3). **d**, Immunoblot of SMAD2, SMAD3, and corresponding phospho-deficient mutants in ERK2^D319N^ expressing MCF10A cells. **e**, Immunoblot of SMAD2, SMAD3, and corresponding phospho-deficient mutants in TGF-β1 (5 ng/ml) stimulated MCF10A cells. **f**, Brightfield images corresponding to samples in **d**. **g**, Brightfield images corresponding to samples in **e**.

**Extended Data Fig. 3.**
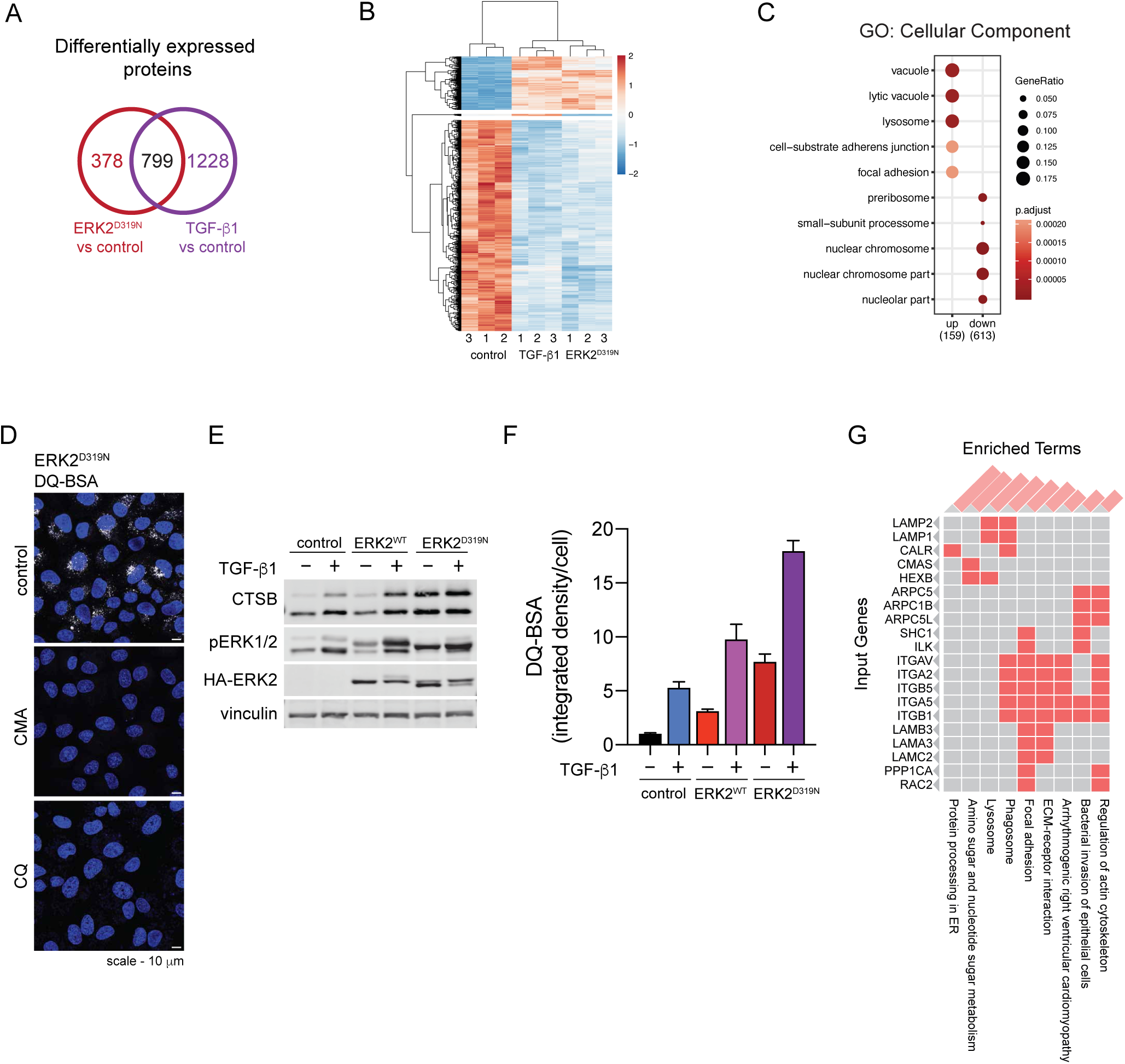
Lysosome induction in EMT. **a**, Venn diagram and **b**, heatmap of differentially expressed proteins in ERK2^D319N^ expressing and TGF-β1 stimulated cells (5 ng/ml). **c**, Enrichment analysis of statistically significant (p>0.001) up and down-regulated proteins in the two models of EMT. **d**, Fluorescence microscopy of DQ-BSA in ERK2^D319N^ expressing cells treated with 100 nM concanamycin A (CMA) or 50 μM chloroquine (CQ). **e-f**, Immunoblot and corresponding DQ-BSA fluorescence integrated density in MCF10A cells expressing ERK2^WT^ or ERK2^D319N^ stimulated with TGF-β1 (5ng/ml) (n = 3). **g**, Enrichment clustergram of significantly increased proteins from proteomics of ERK2^D319N^ expressing cells on day 3. Red bars (top) correspond to the combined score of listed KEGG terms.

**Extended Data Fig. 4.**
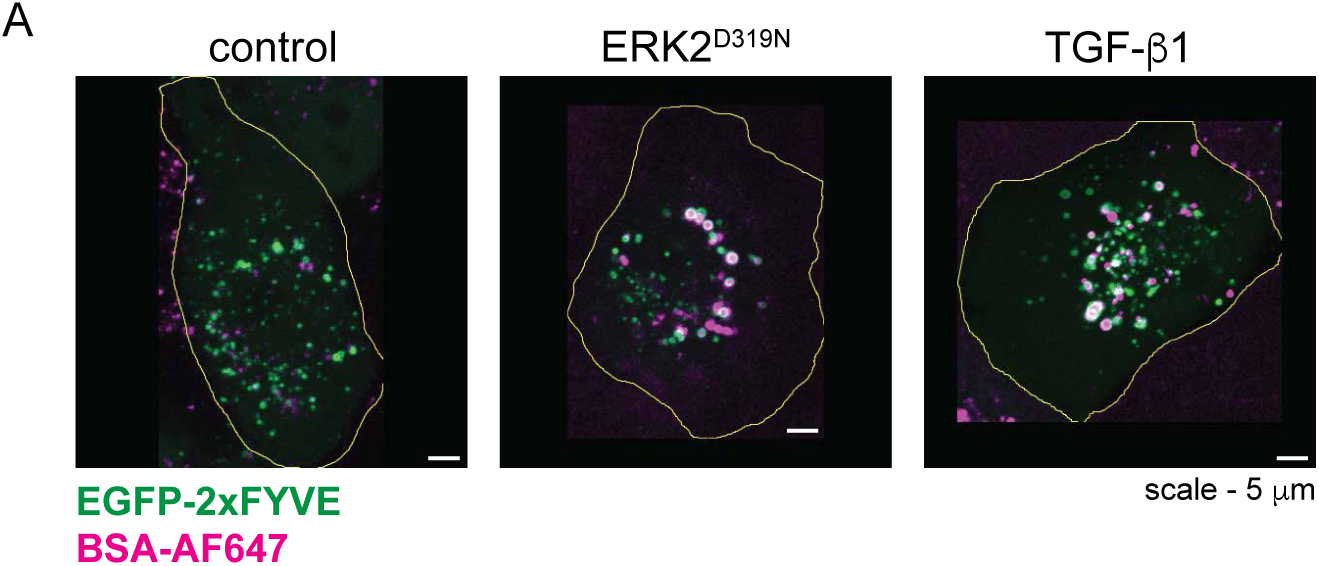
Macropinocytosis in EMT. **a**, Live cell fluorescence microscopy of Alexa Fluor 647 conjugated BSA uptake in MCF10A cells expressing EGFP-2xFYVE.

**Extended Data Fig. 5.**
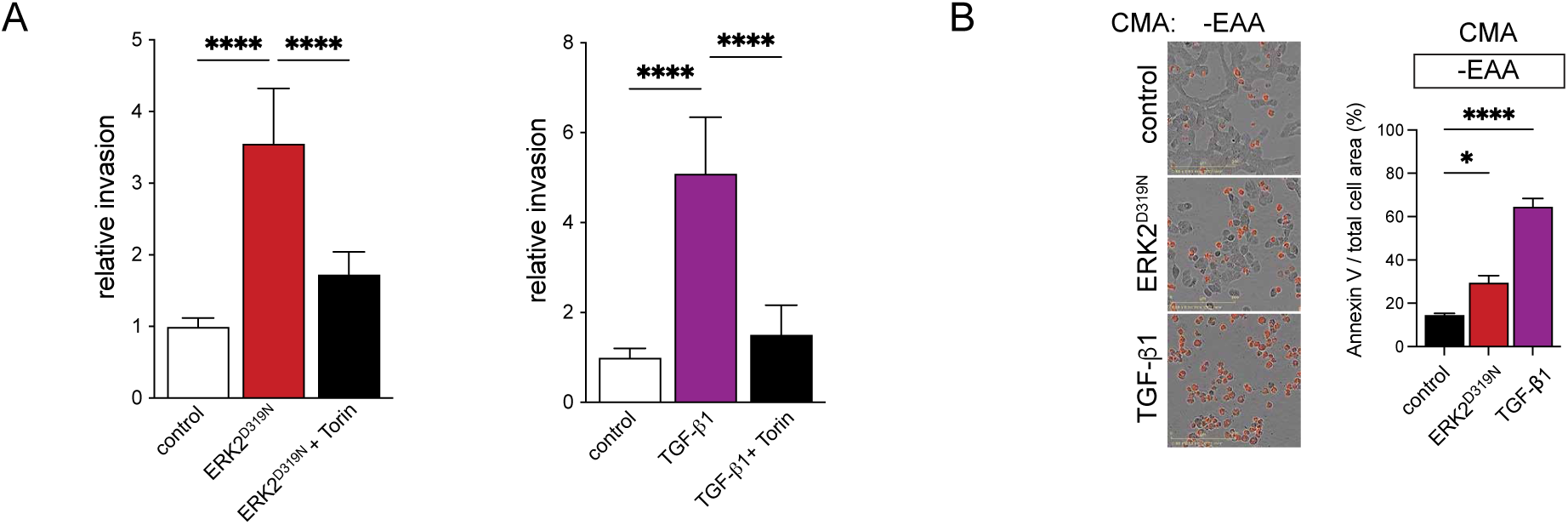
Effects of mTOR-inhibition on invasion and lysosome inhibition on EMT cell toxicity. **a**, Matrigel invasion assay of MCF10A cells expressing ERK2^D319N^ or stimulated with TGF-β1 (5 ng/ml) and treated with 250 nM Torin at the time of seeding in the Boyden chamber. Cell invasion was analyzed after 24 h (n = 3). **b**, Annexin V staining of MCF10A control cells and those undergoing EMT starved of essential AA and glutamine (-EAAQ) for 48 h in the presence of 100 nM concanamycin A (CMA) (n = 3).

**Extended Data Fig. 6.**
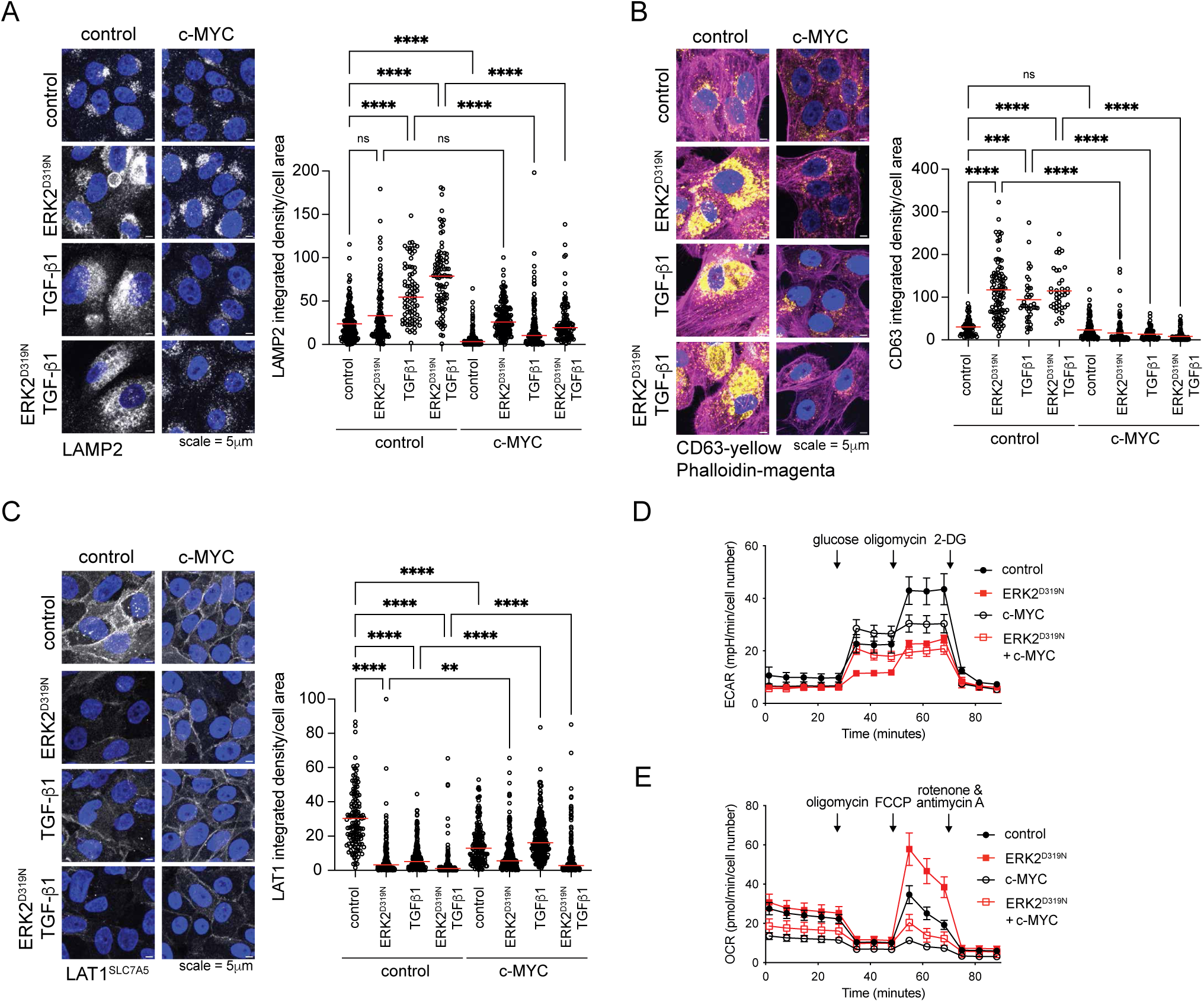
c-MYC opposes EMT associated amino acid transport changes and metabolic switch. **a**, Immunofluorescence of LAMP2 in MCF10A undergoing EMT with c-MYC expression. Data collected over 4-5 fields, n ≥ 23 cells per group. **b**, Immunofluorescence of CD63 and phalloidin in MCF10A undergoing EMT with c-MYC expression. Data collected over 2 fields, n ≥ 36 cells per group. **c**, Immunofluorescence of LAT1/SCL7A5 in MCF10A undergoing EMT with c-MYC expression. Data collected over 3 fields, n ≥ 100 cells per group. **d**, Glycolysis stress test profile indicating changes to the extracellular acidification rate (ECAR) of cells expressing control, ERK2^D319N^, c-MYC or ERK2^D319N^ with c-MYC (Day 3). **e**, Mitochondrial stress test profile indicating changes to the oxygen consumption rate (OCR) of cells expressing control, ERK2^D319N^, c-MYC or ERK2^D319N^ with c-MYC (Day 3).

## References

1. Senft, D. & Ronai, Z.E. Adaptive Stress Responses During Tumor Metastasis and Dormancy. Trends Cancer 2, 429–442 (2016).

2. Shen, M. & Kang, Y. Stresses in the metastatic cascade: molecular mechanisms and therapeutic opportunities. Genes Dev 34, 1577–1598 (2020).

3. Lu, W. & Kang, Y. Epithelial-Mesenchymal Plasticity in Cancer Progression and Metastasis. Dev Cell 49, 361–374 (2019).

4. Thiery, J.P. & Sleeman, J.P. Complex networks orchestrate epithelial-mesenchymal transitions. Nat Rev Mol Cell Biol 7, 131–142 (2006).

5. Zavadil, J. et al. Genetic programs of epithelial cell plasticity directed by transforming growth factor-beta. Proc Natl Acad Sci U S A 98, 6686–6691 (2001).

6. Tam, W.L. et al. Protein kinase C alpha is a central signaling node and therapeutic target for breast cancer stem cells. Cancer Cell 24, 347–364 (2013).

7. Buonato, J.M. & Lazzara, M.J. ERK1/2 blockade prevents epithelial-mesenchymal transition in lung cancer cells and promotes their sensitivity to EGFR inhibition. Cancer Res 74, 309–319 (2014).

8. Su, J. et al. TGF-beta orchestrates fibrogenic and developmental EMTs via the RAS effector RREB1. Nature 577, 566–571 (2020).

9. Shin, S., Dimitri, C.A., Yoon, S.O., Dowdle, W. & Blenis, J. ERK2 but not ERK1 induces epithelial-to-mesenchymal transformation via DEF motif-dependent signaling events. Mol Cell 38, 114–127 (2010).

10. Gomes, A.P. et al. Dynamic Incorporation of Histone H3 Variants into Chromatin Is Essential for Acquisition of Aggressive Traits and Metastatic Colonization. Cancer Cell 36, 402–417 e413 (2019).

11. Shin, S. et al. ERK2 regulates epithelial-to-mesenchymal plasticity through DOCK10-dependent Rac1/FoxO1 activation. Proc Natl Acad Sci U S A 116, 2967–2976 (2019).

12. Kong, X. et al. Cancer drug addiction is relayed by an ERK2-dependent phenotype switch. Nature 550, 270–274 (2017).

13. Gagliardi, M. et al. Differential functions of ERK1 and ERK2 in lung metastasis processes in triple-negative breast cancer. Sci Rep 10, 8537 (2020).

14. Bergers, G. & Fendt, S.M. The metabolism of cancer cells during metastasis. Nat Rev Cancer 21, 162–180 (2021).

15. Ballabio, A. & Bonifacino, J.S. Lysosomes as dynamic regulators of cell and organismal homeostasis. Nat Rev Mol Cell Biol 21, 101–118 (2020).

16. Davidson, S.M. & Vander Heiden, M.G. Critical Functions of the Lysosome in Cancer Biology. Annu Rev Pharmacol Toxicol 57, 481–507 (2017).

17. Palm, W. & Thompson, C.B. Nutrient acquisition strategies of mammalian cells. Nature 546, 234–242 (2017).

18. Finicle, B.T., Jayashankar, V. & Edinger, A.L. Nutrient scavenging in cancer. Nat Rev Cancer 18, 619–633 (2018).

19. Banushi, B., Joseph, S.R., Lum, B., Lee, J.J. & Simpson, F. Endocytosis in cancer and cancer therapy. Nat Rev Cancer 23, 450–473 (2023).

20. Recouvreux, M.V. et al. Glutamine depletion regulates Slug to promote EMT and metastasis in pancreatic cancer. J Exp Med 217 (2020).

21. King, B., Araki, J., Palm, W. & Thompson, C.B. Yap/Taz promote the scavenging of extracellular nutrients through macropinocytosis. Genes Dev 34, 1345–1358 (2020).

22. Marine, J.C., Dawson, S.J. & Dawson, M.A. Non-genetic mechanisms of therapeutic resistance in cancer. Nat Rev Cancer 20, 743–756 (2020).

23. Brenan, L. et al. Phenotypic Characterization of a Comprehensive Set of MAPK1/ERK2 Missense Mutants. Cell Rep 17, 1171–1183 (2016).

24. Padua, D. et al. TGFbeta primes breast tumors for lung metastasis seeding through angiopoietin-like 4. Cell 133, 66–77 (2008).

25. Jin, X. et al. A metastasis map of human cancer cell lines. Nature 588, 331–336 (2020).

26. Lehuede, C., Dupuy, F., Rabinovitch, R., Jones, R.G. & Siegel, P.M. Metabolic Plasticity as a Determinant of Tumor Growth and Metastasis. Cancer Res 76, 5201–5208 (2016).

27. Ravichandran, M. et al. Coordinated Transcriptional and Catabolic Programs Support Iron-Dependent Adaptation to RAS-MAPK Pathway Inhibition in Pancreatic Cancer. Cancer Discov 12, 2198–2219 (2022).

28. Massague, J. & Sheppard, D. TGF-beta signaling in health and disease. Cell 186, 4007–4037 (2023).

29. Xie, L. et al. Activation of the Erk pathway is required for TGF-beta1-induced EMT in vitro. Neoplasia 6, 603–610 (2004).

30. Mizushima, N., Yoshimori, T. & Levine, B. Methods in mammalian autophagy research. Cell 140, 313–326 (2010).

31. N’Diaye, E.N. et al. PLIC proteins or ubiquilins regulate autophagy-dependent cell survival during nutrient starvation. EMBO Rep 10, 173–179 (2009).

32. Verdon, Q. et al. SNAT7 is the primary lysosomal glutamine exporter required for extracellular protein-dependent growth of cancer cells. Proc Natl Acad Sci U S A 114, E3602–E3611 (2017).

33. Zoncu, R. et al. mTORC1 senses lysosomal amino acids through an inside-out mechanism that requires the vacuolar H(+)-ATPase. Science 334, 678–683 (2011).

34. Lamouille, S. & Derynck, R. Cell size and invasion in TGF-beta-induced epithelial to mesenchymal transition is regulated by activation of the mTOR pathway. J Cell Biol 178, 437–451 (2007).

35. Bandyopadhyay, U. et al. Leucine retention in lysosomes is regulated by starvation. Proc Natl Acad Sci U S A 119 (2022).

36. Jayashankar, V. & Edinger, A.L. Macropinocytosis confers resistance to therapies targeting cancer anabolism. Nat Commun 11, 1121 (2020).

37. Chen, C.R., Kang, Y., Siegel, P.M. & Massague, J. E2F4/5 and p107 as Smad cofactors linking the TGFbeta receptor to c-myc repression. Cell 110, 19–32 (2002).

38. Yagi, K. et al. c-myc is a downstream target of the Smad pathway. J Biol Chem 277, 854–861 (2002).

39. Gao, P. et al. c-Myc suppression of miR-23a/b enhances mitochondrial glutaminase expression and glutamine metabolism. Nature 458, 762–765 (2009).

40. Hayashi, K., Jutabha, P., Endou, H. & Anzai, N. c-Myc is crucial for the expression of LAT1 in MIA Paca-2 human pancreatic cancer cells. Oncol Rep 28, 862–866 (2012).

41. Osthus, R.C. et al. Deregulation of glucose transporter 1 and glycolytic gene expression by c-Myc. J Biol Chem 275, 21797–21800 (2000).

42. Wise, D.R. et al. Myc regulates a transcriptional program that stimulates mitochondrial glutaminolysis and leads to glutamine addiction. Proc Natl Acad Sci U S A 105, 18782–18787 (2008).

43. Zhao, X., Petrashen, A.P., Sanders, J.A., Peterson, A.L. & Sedivy, J.M. SLC1A5 glutamine transporter is a target of MYC and mediates reduced mTORC1 signaling and increased fatty acid oxidation in long-lived Myc hypomorphic mice. Aging Cell 18, e12947 (2019).

44. Annunziata, I. et al. MYC competes with MiT/TFE in regulating lysosomal biogenesis and autophagy through an epigenetic rheostat. Nat Commun 10, 3623 (2019).

45. Dejure, F.R. & Eilers, M. MYC and tumor metabolism: chicken and egg. EMBO J 36, 3409–3420 (2017).

46. Liu, H. et al. MYC suppresses cancer metastasis by direct transcriptional silencing of alphav and beta3 integrin subunits. Nat Cell Biol 14, 567–574 (2012).

47. Janda, E. et al. Raf plus TGFbeta-dependent EMT is initiated by endocytosis and lysosomal degradation of E-cadherin. Oncogene 25, 7117–7130 (2006).

48. Kern, U., Wischnewski, V., Biniossek, M.L., Schilling, O. & Reinheckel, T. Lysosomal protein turnover contributes to the acquisition of TGFbeta-1 induced invasive properties of mammary cancer cells. Mol Cancer 14, 39 (2015).

49. Palacios, F., Tushir, J.S., Fujita, Y. & D’Souza-Schorey, C. Lysosomal targeting of E-cadherin: a unique mechanism for the down-regulation of cell-cell adhesion during epithelial to mesenchymal transitions. Mol Cell Biol 25, 389–402 (2005).

50. Jiang, Y., Woosley, A.N., Sivalingam, N., Natarajan, S. & Howe, P.H. Cathepsin-B-mediated cleavage of Disabled-2 regulates TGF-beta-induced autophagy. Nat Cell Biol 18, 851–863 (2016).

51. Mutvei, A.P., Nagiec, M.J. & Blenis, J. Balancing lysosome abundance in health and disease. Nat Cell Biol 25, 1254–1264 (2023).

52. Puertollano, R., Ferguson, S.M., Brugarolas, J. & Ballabio, A. The complex relationship between TFEB transcription factor phosphorylation and subcellular localization. EMBO J 37 (2018).

53. Perera, R.M. et al. Transcriptional control of autophagy-lysosome function drives pancreatic cancer metabolism. Nature 524, 361–365 (2015).

54. Commisso, C. et al. Macropinocytosis of protein is an amino acid supply route in Ras-transformed cells. Nature 497, 633–637 (2013).

55. Palm, W., Araki, J., King, B., DeMatteo, R.G. & Thompson, C.B. Critical role for PI3-kinase in regulating the use of proteins as an amino acid source. Proc Natl Acad Sci U S A 114, E8628–E8636 (2017).

56. Hapke, R.Y. & Haake, S.M. Hypoxia-induced epithelial to mesenchymal transition in cancer. Cancer Lett 487, 10–20 (2020).

57. Shibue, T. & Weinberg, R.A. EMT, CSCs, and drug resistance: the mechanistic link and clinical implications. Nat Rev Clin Oncol 14, 611–629 (2017).

58. Okreglak, V. et al. Cell cycle-linked vacuolar pH dynamics regulate amino acid homeostasis and cell growth. Nat Metab 5, 1803–1819 (2023).

59. Todorova, P.K. et al. Amino acid intake strategies define pluripotent cell states. Nat Metab 6, 127–140 (2024).

60. Gu, Z., Noss, E.H., Hsu, V.W. & Brenner, M.B. Integrins traffic rapidly via circular dorsal ruffles and macropinocytosis during stimulated cell migration. J Cell Biol 193, 61–70 (2011).

61. Zdzalik-Bielecka, D. et al. The GAS6-AXL signaling pathway triggers actin remodeling that drives membrane ruffling, macropinocytosis, and cancer-cell invasion. Proc Natl Acad Sci U S A 118 (2021).

62. Davis, R.T. et al. Transcriptional diversity and bioenergetic shift in human breast cancer metastasis revealed by single-cell RNA sequencing. Nat Cell Biol 22, 310–320 (2020).

63. LeBleu, V.S. et al. PGC-1alpha mediates mitochondrial biogenesis and oxidative phosphorylation in cancer cells to promote metastasis. Nat Cell Biol 16, 992–1003, 1001-1015 (2014).

64. Rainero, E. et al. Ligand-Occupied Integrin Internalization Links Nutrient Signaling to Invasive Migration. Cell Rep 10, 398–413 (2015).

65. Muranen, T. et al. Starved epithelial cells uptake extracellular matrix for survival. Nat Commun 8, 13989 (2017).

66. Jiang, G. et al. Targeted drug delivery system inspired by macropinocytosis. J Control Release 359, 302–314 (2023).

67. Hoogenboezem, E.N. & Duvall, C.L. Harnessing albumin as a carrier for cancer therapies. Adv Drug Deliv Rev 130, 73–89 (2018).

68. Li, R. et al. Therapeutically reprogrammed nutrient signalling enhances nanoparticulate albumin bound drug uptake and efficacy in KRAS-mutant cancer. Nat Nanotechnol 16, 830–839 (2021).

## References

69. Gardner, E.E. et al. Lineage-specific intolerance to oncogenic drivers restricts histological transformation. Science 383, eadj1415 (2024).

70. Fukushima, H. et al. SCF-mediated Cdh1 degradation defines a negative feedback system that coordinates cell-cycle progression. Cell Rep 4, 803–816 (2013).

71. Carvalho, B.S. & Irizarry, R.A. A framework for oligonucleotide microarray preprocessing. Bioinformatics 26, 2363–2367 (2010).

72. Irizarry, R.A. et al. Exploration, normalization, and summaries of high density oligonucleotide array probe level data. Biostatistics 4, 249–264 (2003).

73. Ritchie, M.E. et al. limma powers differential expression analyses for RNA-sequencing and microarray studies. Nucleic Acids Res 43, e47 (2015).

74. Albrecht, M. et al. TTCA: an R package for the identification of differentially expressed genes in time course microarray data. BMC Bioinformatics 18, 33 (2017).

75. Vaga, S. et al. Phosphoproteomic analyses reveal novel cross-modulation mechanisms between two signaling pathways in yeast. Mol Syst Biol 10, 767 (2014).

76. Erickson, B.K. et al. Active Instrument Engagement Combined with a Real-Time Database Search for Improved Performance of Sample Multiplexing Workflows. J Proteome Res 18, 1299–1306 (2019).

77. Eng, J.K., McCormack, A.L. & Yates, J.R. An approach to correlate tandem mass spectral data of peptides with amino acid sequences in a protein database. J Am Soc Mass Spectrom 5, 976–989 (1994).

78. Washburn, M.P. The H-index of ’an approach to correlate tandem mass spectral data of peptides with amino acid sequences in a protein database’. J Am Soc Mass Spectrom 26, 1799–1803 (2015).

79. Elias, J.E. & Gygi, S.P. Target-decoy search strategy for increased confidence in large-scale protein identifications by mass spectrometry. Nat Methods 4, 207–214 (2007).

80. Huttlin, E.L. et al. A tissue-specific atlas of mouse protein phosphorylation and expression. Cell 143, 1174–1189 (2010).

81. Xie, Z. et al. Gene Set Knowledge Discovery with Enrichr. Curr Protoc 1, e90 (2021).

82. Dieterle, F., Ross, A., Schlotterbeck, G. & Senn, H. Probabilistic quotient normalization as robust method to account for dilution of complex biological mixtures. Application in 1H NMR metabonomics. Anal Chem 78, 4281–4290 (2006).

83. Do, K.T. et al. Characterization of missing values in untargeted MS-based metabolomics data and evaluation of missing data handling strategies. Metabolomics 14, 128 (2018).

84. Rueden, C.T. et al. ImageJ2: ImageJ for the next generation of scientific image data. BMC Bioinformatics 18, 529 (2017).

85. Schindelin, J., et al. Fiji: an open-source platform for biological-image analysis. Nat Methods 9, 676-682 (2012).

86. Ito, S. & Karnovsky, M.J. Formaldehyde-glutaraldehyde fixatives containing trinitro compounds. Journal of Cell Biology 39, 168a–169a (1968).

87. de Bruijn, W.C. Glycogen, its chemistry and morphologic appearance in the electron microscope. I. A modified OsO 4 fixative which selectively contrasts glycogen. J Ultrastruct Res 42, 29–50 (1973).

88. Venable, J.H. & Coggeshall, R. A Simplified Lead Citrate Stain for Use in Electron Microscopy. J Cell Biol 25, 407–408 (1965).

